# PyLipID: A Python package for analysis of protein-lipid interactions from MD simulations

**DOI:** 10.1101/2021.07.14.452312

**Authors:** Wanling Song, Robin A. Corey, T. Bertie Ansell, C. Keith Cassidy, Michael R. Horrell, Anna L. Duncan, Phillip J. Stansfeld, Mark S.P. Sansom

**Affiliations:** Department of Biochemistry, University of Oxford, South Parks Road, Oxford, OX1 3QU, UK; Rahko, Clifton House, 46 Clifton Terrace, Finsbury Park, London N4 3JP, UK; School of Life Sciences & Department of Chemistry, University of Warwick, Coventry, CV4 7AL, UK

**Author notes:** Co-corresponding authors &. For submission to JCTC.

## Abstract

Lipids play important modulatory and structural roles for membrane proteins. Molecular dynamics simulations are frequently used to provide insights into the nature of these protein-lipid interactions. Systematic comparative analysis requires tools that provide algorithms for objective assessment of such interactions. We introduce PyLipID, a python package for the identification and characterization of specific lipid interactions and binding sites on membrane proteins from molecular dynamics simulations. PyLipID uses a community analysis approach for binding site detection, calculating lipid residence times for both the individual protein residues and the detected binding sites. To assist structural analysis, PyLipID produces representative bound lipid poses from simulation data, using a density-based scoring function. To estimate residue contacts robustly, PyLipID uses a dual-cutoff scheme to differentiate between lipid conformational rearrangements whilst bound from full dissociation events. In addition to the characterization of protein-lipid interactions, PyLipID is applicable to analysis of the interactions of membrane proteins with other ligands. By combining automated analysis, efficient algorithms, and open-source distribution, PyLipID facilitates the systematic analysis of lipid interactions from large simulation datasets of multiple species of membrane proteins.

**ToC Graphic:** 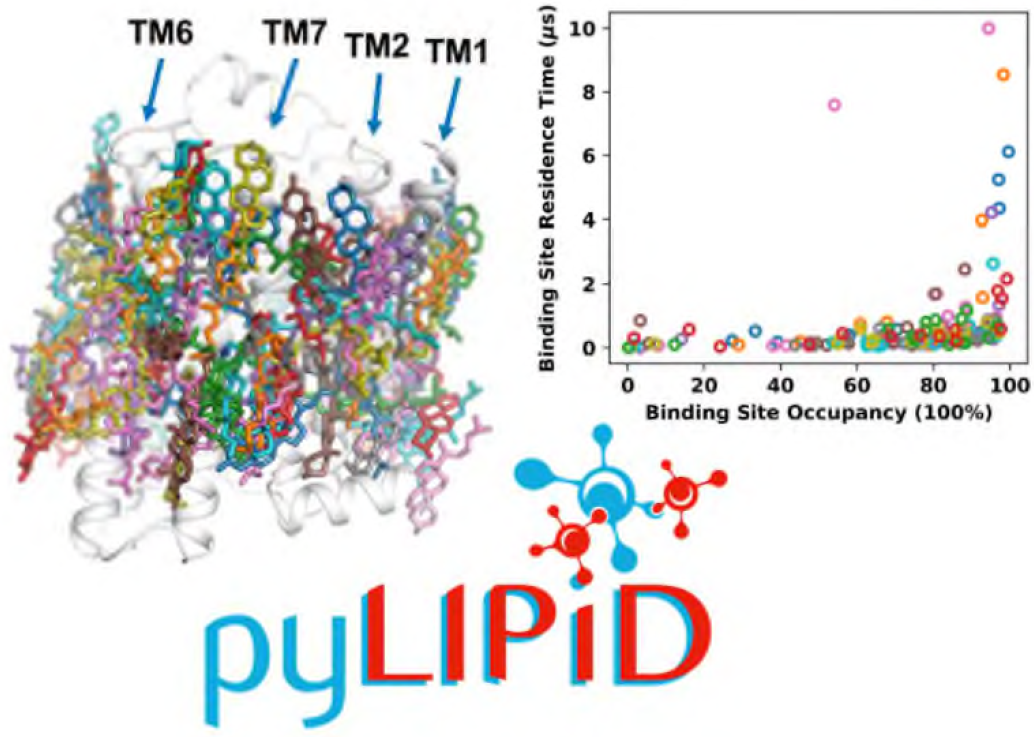

## INTRODUCTION

Cell membranes typically contain hundreds of different lipid species, asymmetrically distributed between two membrane leaflets ^1-2^. These lipid molecules are locally organized into lateral domains of distinct composition ^3-4^. The combination of these various chemical structures and microdomains results in a diverse lipid landscape that is fully exploited by membrane proteins, especially those involved in cellular signaling ^5-6^. The regulatory roles of membrane lipids include ion channel activation ^7-8^ and allosteric modulation of GPCRs and other receptors ^9-13^. Lipid molecules may also strengthen domain and/or subunit interactions in more complex membrane proteins ^14-15^. It is therefore of importance to characterize protein-lipid interactions in order to reach an understanding of the dynamics and functions of membrane proteins.

A number of biophysical techniques can reveal the presence of protein-lipid interactions e.g. ^16-17^. In particular, following recent advances in single particle cryo-EM ^18^ including the use of nanodiscs to preserve a lipid bilayer environment ^19^, increasing numbers of membrane proteins structures have been determined at near atomic resolution with bound lipids present in the structures ^20^. These membrane protein structures provide gateways for understanding how lipids may modulate protein function but also pose challenges regarding the identification of interacting lipid species.

Computational approaches, especially molecular dynamics (MD) simulations, have played an increasingly important role as a high throughput “computational microscope” ^21^ for the identification of protein-lipid interactions. Thanks to ongoing increases in computer power, development of improved atomistic and coarse-grained force fields ^22^, and development of tools to automate setup of membrane simulations ^23-24^, MD simulations have been applied to many membrane proteins and lipids, providing invaluable structural and mechanistic insights into their protein-lipid interactions ^25-27^.

For the study of protein-lipid interactions, CG force fields can explore lipid binding sites in an unbiased fashion with sufficient sampling due to the decreased degrees of freedom of the underlying model. The Martini forcefield ^28-30^ is widely used for biomembrane applications. Simulations using Martini have identified lipid binding sites on a range of proteins ^31-33^, and have assisted the interpretation of lipid-like density in cryo-EM maps^34^. Some simulation studies have adopted a serial multiscale approach in which CG simulations are used to probe the lipid binding sites and bound lipid identities followed by atomistic simulations to study residue-level protein-lipid interactions. The conversion of Martini models to atomistic ones can be achieved by resolution back-mapping tools ^35-38^.

With the increasing number of membrane protein structures determined at high resolution by cryo-EM and the increasing complexity of simulated membranes, the use of MD simulations to study protein-lipid interactions is accompanied by two challenges:

(1) How to automatically determine lipid binding sites from simulations? Some simulation studies have used the average lipid density to approximately locate lipid binding sites and subsequently manually assigned bound poses. Such an approach includes an element of subjectivity and may be a bottleneck for large scale comparative simulations. So, can we determine the lipid binding sites automatically via a statistically robust method? Additionally, can we systematically produce representative bound poses from the trajectories for further analysis?

(2) How to optimally quantify and characterize lipid interactions? Simulation studies have used e.g. average lipid occupancies or the fraction of trajectory frames in which lipid contacts are formed to a given residue to measure lipid interactions. Can we rigorously calculate lipid interactions with binding sites in addition to individual residues to allow for more direct comparison with experiments?

To provide a unified solution to the above-mentioned problems, we have developed a Python package, PyLipID, to assist analysis of protein-lipid interactions from MD simulations. PyLipID identifies binding sites by calculating the community structures in the interaction network of protein residues. This method was initially applied to the analysis of cholesterol ^39^ and other lipid ^33^ binding sites on the Kir2.2 channel. Based on the identified binding sites, PyLipID can find representative bound poses for each site. This is achieved by evaluating all the bound poses using an empirical scoring function of the lipid density in the chosen binding site. This functionality allows for further structural analysis of the protein-lipid interactions and makes it possible to automate pipelines for converting bound lipids poses in CG models into atomistic ones for use in multiscale simulation studies. PyLipID can also cluster the bound poses for binding sites to provide a more in-depth analysis of the lipid interactions. To describe lipid interactions, PyLipID calculates residence times, in addition to other commonly used metrics such as averaged interaction duration, lipid occupancy, and the average number of surrounding lipids, for both individual protein residues and the calculated binding sites. The calculation of residence times reveals the dynamical behavior of bound lipids, and calculations based on binding sites allow for improved characterization of the binding events. Notably, PyLipID uses a dual-cutoff scheme to deal with the ‘rattling in a cage’ effect sometimes seen in protein-lipid simulations.

In the following sections, we first introduce the methodological details of PyLipID. Then, we illustrate the PyLipID analysis pipeline using cholesterol binding to a panel of G-protein coupled receptors (GPCRs) as an example. Subsequently, we present two cases of the application of PyLipID to interactions of membrane proteins with phospholipids illustrating the potential application of PyLipID to assist the interpretation of lipid-like densities in cryo-EM maps. Finally, we demonstrate the application of PyLipID to non-lipid molecules, using it characterize ethanol binding to the cytoplasmic domain of the *B. subtilis* McpB chemoreceptor as seen in atomistic simulations ^40^.

## METHODS

PyLipID is an open-source package available on GitHub (https://github.com/wlsong/PyLipID). The documentation and tutorials can be found at the ReadtheDocs server https://pylipid.readthedocs.io. A tutorial script that runs the PyLipID analysis can be found at the documentation website.

### Overview of code

The current PyLipID package contains four modules: `api`, `func`, `plot` and `util`. The `api` is the outer layer module that handles the analysis workflow and provides some convenient functions for plotting and saving data (Fig. 1), whereas the remaining modules provide functions that are deployed by `api` for the heavy lifting in the analysis (SI Fig. S1). Such a structure allows for extension of PyLipID functionalities with minimal changes to the code base. PyLipID reports results in various forms.

**Figure 1.**
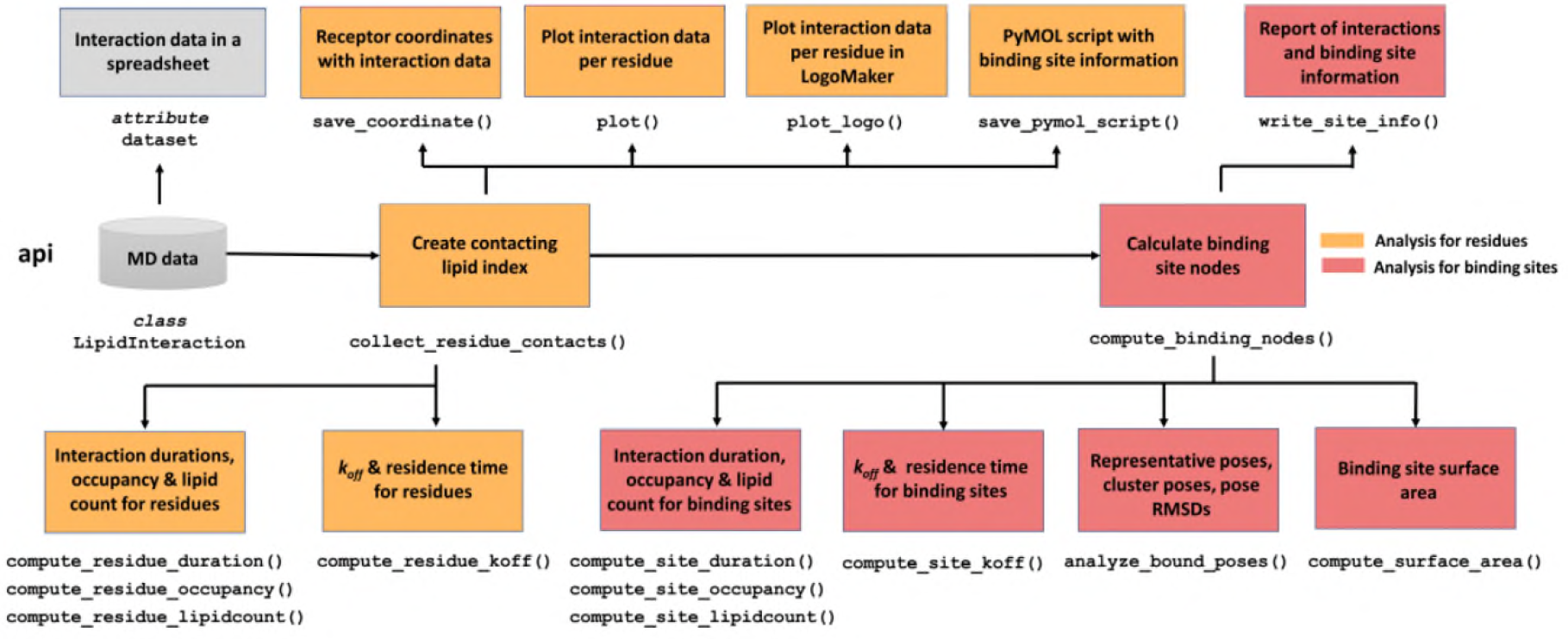
PyLipID package design: api module structure. `api` is the outer layer module and its main class `LipidInteraction` handles the analysis workflow. The class object `LipidInteraction` loads the trajectory data, and the methods of this class object carries out the analysis for protein residues (yellow boxes) and for binding sites (red boxes). This class object also has an attribute `dataset`, which is a spreadsheet object storing interaction data and allows for further manipulation. PyLipID has another 3 modules, `func`, `plot` and `util`, which provides functions for doing the computationally intensive analysis, as used by `LipidInteraction` (SI Fig S1).

*api*. This module contains the main python class `LipidInteraction` that reads trajectory information, analyzes lipid interactions and writes/plots interaction data. The PyLipID analyses are carried out by the class methods of `LipidInteraction`, which can be divided into two groups: methods for analysis of interactions with protein residues and with the calculated binding sites. Each group has a core function to collect/calculate the required data for the rest of the functions in that segment, *i*.*e*. `collect_residue_contacts()`that builds lipid index for residues as a function of time for residue analysis; and `compute_binding_nodes()` that calculates the binding sites using the interaction correlation matrix of the residues. The remainder of the methods in each group are independent of each other and calculate different properties of lipid interactions and of binding site. `LipidInteraction` also has an attribute `dataset` which stores the calculated interaction data in a spreadsheet as a `pandas.DataFrame`, and updates automatically after each of the calculations. It records interaction data for protein residues by row, including interaction residence times, averaged durations, occupancy, and lipid count, and the associated interaction data for the binding site to which the residue belongs. This `pandas.DataFrame` data structure allows for convenient checking of the interaction data and provides users with maximum flexibility to further process PyLipID outputs. For the computationally intensive functions, e.g., calculation of *k*_*off*_, bound poses or binding site surface areas, PyLipID uses a python multiprocessing library to speed up the calculations. Users can specify the number of CPUs these functions can use, otherwise all available CPUs will be used by default.

*func*. This module comprises the following four submodules: `interaction` that contains functions for calculation of continuous contacts using a double cutoff scheme; `kinetics` for calculation of *k*_*off*_ and residence time; `binding_site` for calculation of binding sites using the Louvain method^41^ as well as the analysis of bound poses and surface area; and `clusterer` for clustering the bound poses.

*plot*. This module provides convenient functions to visualize the interaction data, *e*.*g*. plots of *k*_*off*_, interaction as a function of residue index, the correlation matrix of lipid interactions for residues and binding site data.

*util*. This is the location of house-keeping functions. For example, `trajectory` contains functions for obtaining topology information from trajectories.

### Technical features

PyLipID is written in python and compatible with versions 3.6+. It uses MDtraj ^42^ to handle trajectories and coordinates, and thus it is compatible with all major simulation packages. PyLipID reads the molecule topology from trajectories and uses a distance-based method to measure contacts, it is therefore applicable to the calculation of binding characteristics for any type of molecule. In the following section, we will introduce the technical features of PyLipID and their implementation in the code.

#### Lipid topology

The lipid topology information is read from trajectories and contacts are calculated based on the minimum distance of the lipid molecule to the protein. A lipid molecule is considered as being in contact with a residue when the distance of any atoms of the lipid molecule to any atoms of the residue is smaller than the provided distance cutoff. PyLipID also allows for selection of lipid atoms used for defining contacts. This option can be useful for cases in which excluding some atoms (*e*.*g*. the tails of phospholipids) could generate improved definition of binding sites. Given how lipid contact is calculated, PyLipID does not need to store or define any lipid topology information in the code, which allows PyLipID to calculate the contact of any kind of object with a protein based on their distances.

#### Dual-cutoff scheme

Due to the smoothened potentials and/or shallow binding pockets, CG simulations may show a ‘rattling in a cage’ effect, in which lipid molecules undergo rapid changes in protein contacts without full dissociation from a given site, such that the minimum distances between the two contacting objects may experience sudden jumps. This has also been seen in e.g. atomistic simulations of loosely bound cholesterol molecules ^43^. For bound cholesterol molecules in CG simulations, the minimum distance can go up to 0.6-1.0 nm, overlapping with that for non-contacting cholesterols (SI Fig. S2A). This effect can also occur in the atomistic simulations (SI Fig S2B). Using a single distance cutoff to determine the bound/unbound status could thus include unwanted noise.

To deal with these frequently encountered rapid fluctuations in bound pose, PyLipID adopts a dual-cutoff scheme, which uses a lower and upper distance cutoff to measure the status of contact. The duration of a continuous contact is determined from the timepoint when a molecule moves closer than the lower distance cutoff until the timepoint when the molecule moves beyond the upper cutoff distance. The SI provides a more detailed discussion of cut-off values and their impact on binding site calculations (SI text and SI Figures S3-S9). In addition to the contact duration, PyLipID provides another three metrics for characterization of lipid contacts: lipid duration, which is the average duration of the collected contacts; lipid occupancy, which is the percentage of frames in which any lipid contact is formed; and lipid count, which is the number of surrounding molecules of the specified lipid species. Both lipid occupancy and lipid count are calculated using the lower distance cutoff.

#### Residence time

The residence time provides useful insights ^44^ into the dynamic behavior of bound lipids which, due to their interaction with the protein, are no longer diffusive ^31, 33^. Indeed, both prolonged interactions and transient contacts are observed for lipids on the protein surface. The residence time, which is calculated from a survival time correlation function, describes the relaxation of the bound lipids and can be divided into long and short decay periods, which correspond to specific interactions and transient contacts respectively. PyLipID calculates the survival time correlation function *σ*(*t*) as follow:

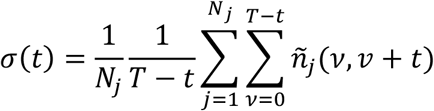

where *T* is the length of the simulation trajectory, *N*_*j*_is the total number of lipid contacts and 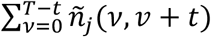 is a binary function that takes the value 1 if the contact of lipid *j* lasts from time *v* to time *v*+*t* and 0 otherwise. The values of *σ*(*t*)are calculated for every value of *t* from 0 to *T*ns, for each time step of the trajectories, and normalized by dividing by σ(0), so that the survival time-correlation function has value 1 at *t* = 0. The normalized survival function is then fitted to a bi-exponential to model the long and short decays of lipid relaxation respectively:

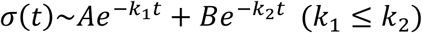

PyLipID takes *k*_l_ as the dissociation constant, *k*_*off*_, and calculates the residence time from *τ* = 1/*k*_*off*_ . It should be noted that providing PyLipID with multiple trajectories of varying length could impact the accuracy of the *k*_*off*_ calculation. PyLipID measures the *r*^2^ of the biexponential fitting to the survival function to show the quality of the *k*_*off*_/residence time estimation. In addition, PyLipID bootstraps the contact durations and measures the *k*_*off*_/residence time of the bootstrapped data, to report how well lipid contacts are sampled from simulations. The lipid contact sampling, the curve-fitting and the bootstrap results can be conveniently checked for individual residues and the calculated binding sites via the *k*_*off*_ plots generated by PyLipID (see Fig. 2 and discussion below for further details).

**Figure 2.**
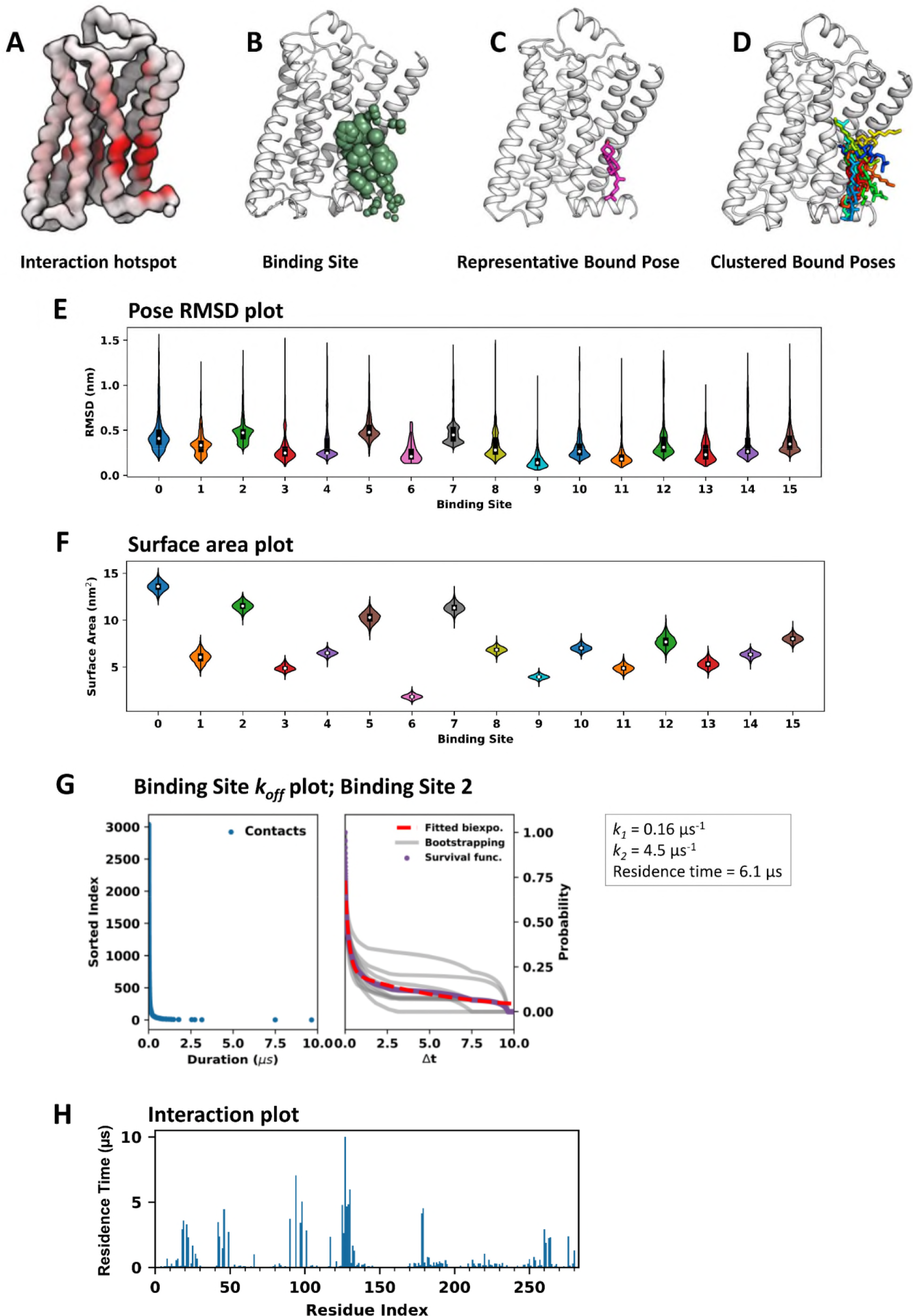
Illustration of PyLipID analysis outputs, using simulations of the β_2_AdR in the presence of cholesterol as an example. PyLipID can save interaction data in the B-factor column of a PDB file of the protein coordinates using `save_coordinate()`. Such a coordinate file can be loaded into a visualization software and colored based on B-factor to show the interaction hotspot (A). PyLipID can generate a python script that maps the binding site information to receptor structure in a PyMOL session, in which residues from the same binding site are shown in spheres in the same color and the sphere scales correspond to their interaction with the lipid. This is accomplished by `save_pymol_script()` (B). The method of `analyze_bound_pose()` can find the representative bound pose for a binding site (C), and cluster all the bound poses in a binding site (D). This method can also calculate the RMSDs of the bound poses for a binding site and provide a convenient plot of the RMSDs (E). The method `compute_surface_area()` calculates binding site surface area as a function of time and plots the surface area data (F). PyLipID calculates interaction residence times for residues using `compute_residue_koff()` and for binding sites using `compute_site_koff()`. Both methods generate k_off_ plots, in which the durations of the collected contacts are plotted in a sorted order in the left panel and the normalized survival function together with the fitted data are plotted in the right panel (G). The `plot()` method can draw the interaction data as a function of residue index (H).

#### Calculation of binding sites

Binding sites are defined based on a community analysis of protein residue-interaction networks that are created from the lipid interaction correlation matrix. Given the basic definition of a lipid binding site, namely a cluster of residues that bind to the same lipid molecule at the same time, PyLipID creates a distance vector that records the distances to all lipid molecules as a function of time for each residue, and constructs a lipid interaction network in which the nodes are the protein residues and the weights are the Pearson correlation coefficients of pairs of residues that are calculated from their distance vectors (SI Fig. S10). PyLipID then decomposes this interaction network into sub-units or communities, which are groups of nodes that are more densely connected internally than with the rest of the network. For the calculation of communities, PyLipID uses the Louvain algorithm ^41^ that finds high modularity network partitions effectively. Modularity, which measures the quality of network partitions, is defined as ^45^

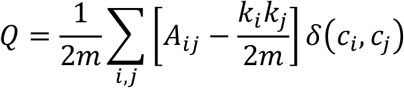

where *A*_*ij*_is the weight of the edge between node *i* and node *j*; *k*_*i*_ is the sum of weights of the nodes attached to the node *i, i*.*e*. the degree of node; *c*_*i*_ is the community to which node *i* is assigned; *δ*(*c*, *c*)is 1 if *i* = *j*and 0 otherwise; and 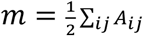, *i*.*e*. the number edges. In the modularity optimization, the Louvain algorithm orders the nodes in the network, and then, one by one, removes and inserts each node in a different community *c*_*i*_ until no significant increase in modularity. After modularity optimization, all the nodes that belong to the same community are merged into a single node, of which the edge weights are the sum of the weights of the comprising nodes. This optimization-aggregation loop is iterated until all nodes are collapsed into one. PyLipID allows for filtering of the communities based on their sizes, *i*.*e*. filtering the binding sites based on the number of comprising residues. By default, PyLipID returns binding sites of at least 4 residues. This filtering step is particularly helpful for analysis of a small number of trajectory frames, in which false correlation is more likely to happen among 2 or 3 residues. The output from this calculation is a list of binding sites containing sets of binding site residue indices.

#### Calculation of representative bound poses

PyLipID evaluates bound poses using an empirical density-based scoring function and writes out the most sampled bound poses for each binding site. The scoring function of a lipid pose at a binding site is defined as:

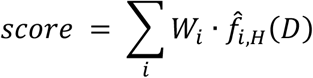

where *W*_*i*_ is the weight given to atom *i* of the lipid molecule, *H* is the bandwidth, and 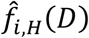 is a multivariate kernel density estimation of the position of atom *i* based on the positions of all bound lipid poses in that binding site. The position of atom *i* is a *p*-variant vector, [*D*_*i*1_, *D*_*i*2_, …, *D*_*ip*_] where *D*_*ip*_ is the minimum distance to the residue *p* of the binding site. PyLipID uses the Gaussian kernel function and, by default, a bandwidth of 0.15. The multivariant kernel density estimation is implemented by *statsmodels* ^46^. Higher weights can be given to *e*.*g*. the headgroup atoms of phospholipids, to generate better defined binding poses, but all lipid atoms are weighted equally by default. In the density estimation, PyLipID uses the relative positions of lipid atoms in the binding site, which makes the analysis of a binding site independent of local protein conformational changes. Lipid poses with the highest scores are considered as the representative bound poses for their binding site and can be written out, along with the protein conformation to which it binds, in any format supported by MDTraj (e.g., pdb and gro). See SI Text for more detailed discussion on the choice of cut-off values and representative bound poses/clustered poses.

#### Clustering of bound lipid poses

PyLipID can cluster the bound lipid poses of a binding site into a user-specified number of clusters using KMeans, in a ‘supervised’ fashion or cluster the poses using a density-based cluster, DBSCAN, in an ‘unsupervised’ fashion. In the former case, the KMeans function from *scikit-learn* ^47^ is used to separate the samples into *n* clusters of equal variances, via minimizing the *inertia*, which is defined as:

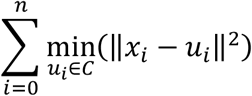

where *u*_*i*_ is the ‘centroid’ of cluster *i*. KMeans scales well with large dataset but performs poorly with clusters of varying sizes and density, which are often the case for lipid poses in a binding site.

When the number of clusters is not provided by user, PyLipID uses the DBSCAN algorithm implemented in *scikit-learn* to find clusters of core samples of high density. A sample point *p* is a core sample if at least *min_samples* points are within distance *ε* of it. A cluster is defined as a set of sample points that are mutually density-connected and density-reachable, *i*.*e*. there is a path ⟨ *p*_1_, *p*_2_, …, *p*_*n*_ ⟩ where each _*i* 1_ is within distance *ε* of _*i*_ for any two *p* in the set. The values of *min_samples* and *ε* determine the performance of this cluster. PyLipID sets the *ε* as the knee point of the *k*-distance graph. Once *ε* is set, the clustering results with all possible *min_samples* are checked using the Silhouette coefficient:

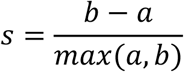

where *a* is the mean distance between a sample and all other points in the same cluster, and *b* is the mean distance between a sample and all other points in the next nearest cluster. The Silhouette coefficient is between -1 and 1, and higher scores suggest better defined clusters. The clustering result with the highest Silhouette score is returned as the optimal clustering results. For writing out the cluster poses, PyLipID randomly selects one pose from each cluster in the case of using KMeans or one from the core samples of each cluster when DBSCAN is used, and writes the selected lipid pose with the protein conformation to which it binds using MDTraj. The relative position of lipid poses in the binding site, *i*.*e*. [*D*_1_, *D*_2_, …, *D*_*i*_] where *D*_*i*_ is the distance vector of atom *i* to the residues in the binding site, is used as the pose coordinates for clustering. Principal component analysis is used to decrease the lipid coordinate dimension before the clustering.

#### Calculation of pose RMSD

The root mean square deviation (RMSD) of a lipid bound pose in a binding site is calculated from the relative position of the pose in the binding site compared to the average position of the bound poses. Thus, the pose RMSD is defined as:

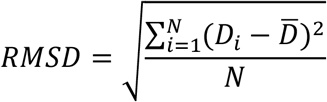

where *D*_*i*_ is the distance vector of atom *i* to the residues in the binding site, 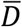 is the average of the distance vectors of atom *i* from all bound poses in the binding site and *N* is the number of atoms in the lipid molecule.

#### Calculation of binding site surface area

The accessible surface area is calculated using the Shrake-Rupley algorithm ^48^. PyLipID strips the protein coordinates out of the simulation system and obtains the accessible surface area of a binding site by summing those of its comprising residues. The surface areas of protein residues are calculated by the shrake_rupley function of MDTraj.

## RESULTS

### PyLipID analysis outputs, illustrated for CG simulations of the interactions of the β_2_AdR with cholesterol

Before describing in detail application cases of PyLipID, we provide a brief overview of PyLipID analysis and outputs (Fig. 2). As an example, we use cholesterol interaction with the β_2_AdR (a GPCR). A more detailed account of GPCR/cholesterol interactions is provided in a subsequent section. We carried out PyLipID analysis using simulation data from 3 repeats. Therefore, the reported durations, occupancies, and lipid counts, for both residues and binding sites, by PyLipID were averaged over the repeats and the residence times were calculated from the durations of lipid contacts collected from all repeats. We also recommend evaluating the impact of different dual cut-offs on binding sites and interaction durations, prior to using PyLipID, to find the optimal values (see SI Text and Figs. S4-S9). For the case of analysis of cholesterol interactions with GPCRs, we chose to use 0.475 and 0.80 nm for the dual cut-offs.

PyLipID outputs results in different forms to assist different analyses. Each analysis is carried out by a method of the class `LipidInteraction`. Users may select specific analysis to implement or use the demo script provided on PyLipID website to run all the analysis once. We first calculated cholesterol interaction, i.e. interaction residence times in this case, with receptor residues via the method `compute_residue_koff()`. To visualize the residue-wise interactions, we used the method `save_coordinate()` to generate a protein databank (PDB) file of the receptor coordinates in which the interaction data are saved in the B factor column, enabling us to check the locations of interaction hotspots (Fig. 2A).

We then calculated the binding sites using the method `compute_binding_nodes()`. After this step, the cholesterol interactions, i.e. residence times in this case, with these binding sites were calculated using `compute_site_koff()`. To assist the visualization of these binding sites, we used the method `save_pymol_script()` to generate a python script that maps the *binding site* information to receptor structure in a PyMOL session, in which residues from the same binding site are shown as spheres in the same color and the sphere scales correspond to their interactions with the lipid (Fig. 2B). This binding site visualization, combined with a binding site summary that was generated by `write_site_info()`,helped to filter through binding sites and find ones of interest. To analyze the structural details of cholesterol interactions, we used `analyze_bound_pose()`to find the representative bound pose for a given binding site (Fig. 2C) and to cluster all the bound poses in a binding site (Fig 2D). In addition, we also calculated other properties of the binding sites/bound poses, including the RMSDs of bound poses via `analyze_bound_poses()` (Fig 2E) and the surface areas of the binding site via `compute_surface_area()` (Fig 2F).

Importantly, when calculating the residence times using either `compute_residue_koff()` or `compute_site_koff()`, PyLipID can also generate the *k*_*off*_ plots, in which the durations of the collected contacts are plotted in a sorted order along with the normalized survival function, fitted bi-exponential curve, and bootstrapped data (Fig. 2G). The quality of the sampling of binding events, which can be checked by the bootstrapping data, and the quality of the evaluation of residence times, which can be checked by *r*^*2*^ of the curve fitting, were checked when we filtered the binding sites.

### Comparative analysis of cholesterol binding sites on selected class A and B GPCRs

The application of PyLipID through python scripts allows for a high-throughput and systematic analysis of large protein-lipid interaction datasets. Here we demonstrate how PyLipID, in conjunction with CG MD simulations, was used to characterize cholesterol binding sites on GPCRs. We performed 3 × 10 µs CG simulations for each of ten species of GPCR (see SI Text and Table S1 for simulation details), embedded within a membrane containing 35% cholesterol and applied PyLipID analysis to study the cholesterol interactions with these receptors.

Combining the residence time profiles and molecular visualization, we found that cholesterol interactions were consistently found between transmembrane helices. However, the strength of the interactions (measured as residence times) varied depending on the receptor and the inter-helical location. We saw stronger cholesterol interactions with β_2_-AdR and D3R at locations around, e.g., TM1, TM7, TM3 and TM4, whereas much weaker interactions were seen in e.g. C-C chemokine receptors and P2Y1 (SI Fig 2-3), suggesting the affinities for cholesterol may varies among receptors and sites.

We then analyzed the lipid bound poses in the binding sites. On average, 14-17 cholesterol binding sites were revealed per receptor and, in total, 153 cholesterol binding sites from the 10 receptors. Aligning the representative bound poses from the 10 tested receptors to the β_2_-AdR structure revealed that cholesterol molecules can be found in most of the inter-helical spaces (Fig. 3A). This is in agreement with a recent analysis of the locations of bound cholesterols in GPCR structures, which reports that cholesterol binding sites lack consensus motifs ^49^. This also lends support to the suggested wedge-like role of cholesterols in stabilizing GPCR conformations ^50^ .

**Figure 3.**
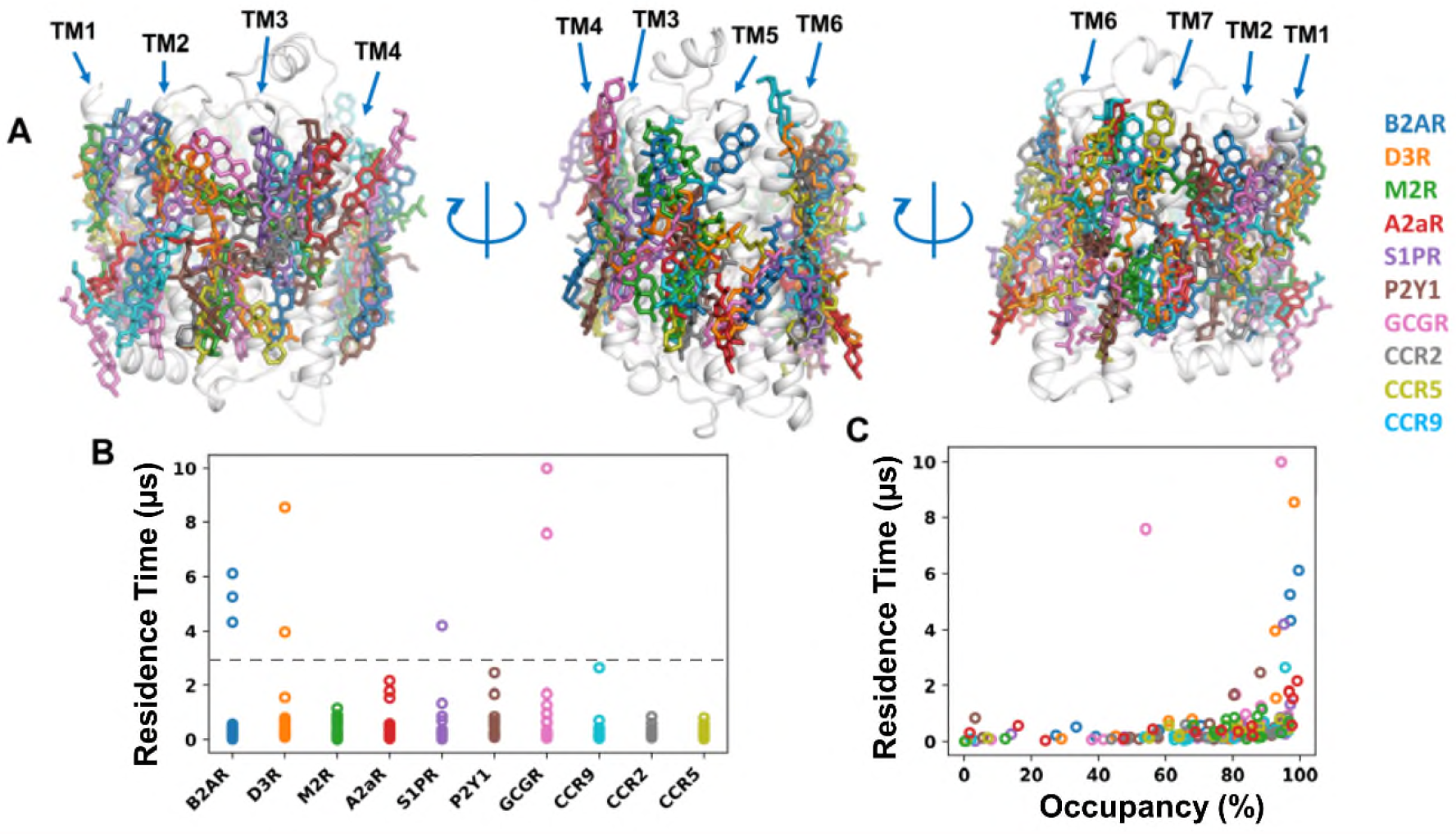
Cholesterol binding sites on GPCRs. (A) The representative cholesterol bound poses of all the binding sites on the 10 GPCRs. The binding sites are aligned to the B2AR structure. (B) Binding site residence times and (C) binding site occupancy calculated from the 10 GPCRs.

We next calculated the binding site residence times and cholesterol occupancies. Most of the cholesterol binding sites had interaction residence times < 3 µs (Fig. 3B). For these sites there was little, if any, correlation between residence time and occupancy (Fig. 3C). The high frequency of cholesterol binding and the relatively short residence times suggest that these cholesterol molecules act as annular lipids around GPCRs, forming a cholesterol solvation shell. However, we also detected a number of binding sites that with residence times > 3 µs (on B2AR, D3R, S1PR and GCGR). With one exception these all had an occupancy of > 90% (Fig. 3C). This suggests that at these sites cholesterol can form longer and more specific interactions.

We then set out to analyze whether there are sequence or structural motifs that determine the length of interaction residence times (i.e. the strength of cholesterol interactions). We first checked whether the size of the binding site affects the interaction. We calculated the binding site surface areas and the buried area, i.e. contacting surface area of bound cholesterols with the receptor. The stronger cholesterol binding sites (i.e. those with residence time > 3 µs) have mid-range sizes, with surface areas between 5-12 nm^2^ (Fig 4A). Visual inspection revealed that the larger binding sites on GPCRs were often flat, shallow, and featureless. The calculation of the buried surface area of cholesterols in the binding sites showed a similar picture, and the bound cholesterols could clearly be separated into two groups (Fig. 4B). For the weaker (non-specific or annular) sites there was perhaps a weak correlation between residence time and buried surface area. For the stronger (specific) sites the contacting surface area did not correlate with the residence times. This suggests that specific binding is more subtly determined than simply the area of the cholesterol binding site on a GPCRs.

**Figure 4.**
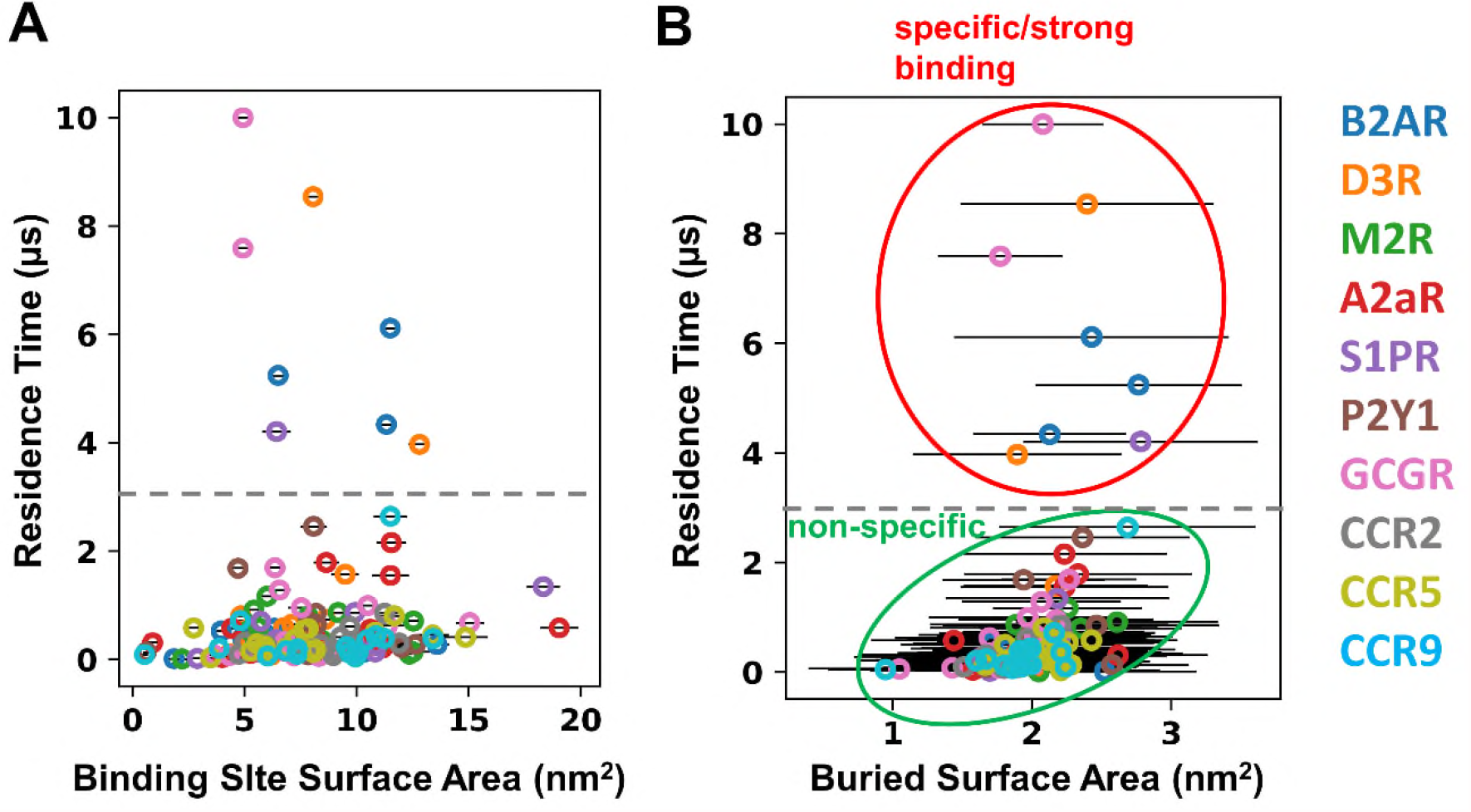
Geometry of cholesterol binding sites on GPCRs. (A) Binding site surface area and (B) the buried surface area of the cholesterol bound in the binding sites on GPCRs. The 3 µs residence time cutoff used to separate non-specific/annular from specific/tight binding interactions is shown as a grey broken line, and the latter two classes are indicated by the green and red ellipses respectively in B.

To explore this further we analyzed the amino acid residue composition of the cholesterol binding sites, looking to see whether longer residence times resulted from a specific composition of the binding sites. We again set a residence time cut-off of 3 µs to separate weaker and strong binding sites, selected from a plot of the sorted residence times (SI Fig S13). We calculated the amino acid composition for each binding site in the two classes (Fig. 5AB). To compare the two sets of data, which have very different sizes, we bootstrapped the data from the nonspecific binding sites. Here, we randomly selected 8 binding sites, and compared their average amino acid composition to the 8 strong binding sites. This comparison revealed that the strong cholesterol binding sites have increased occurrence of Leu, Ala, and Gly residues (Fig 5C). This is broadly consistent with a recent structural analysis ^49^ which failed to reveal distinct sequence motifs for cholesterol binding to GPCRs, but which reported cholesterol microenvironments enriched in Leu, Ala, Ile, and Val residues.

**Figure 5.**
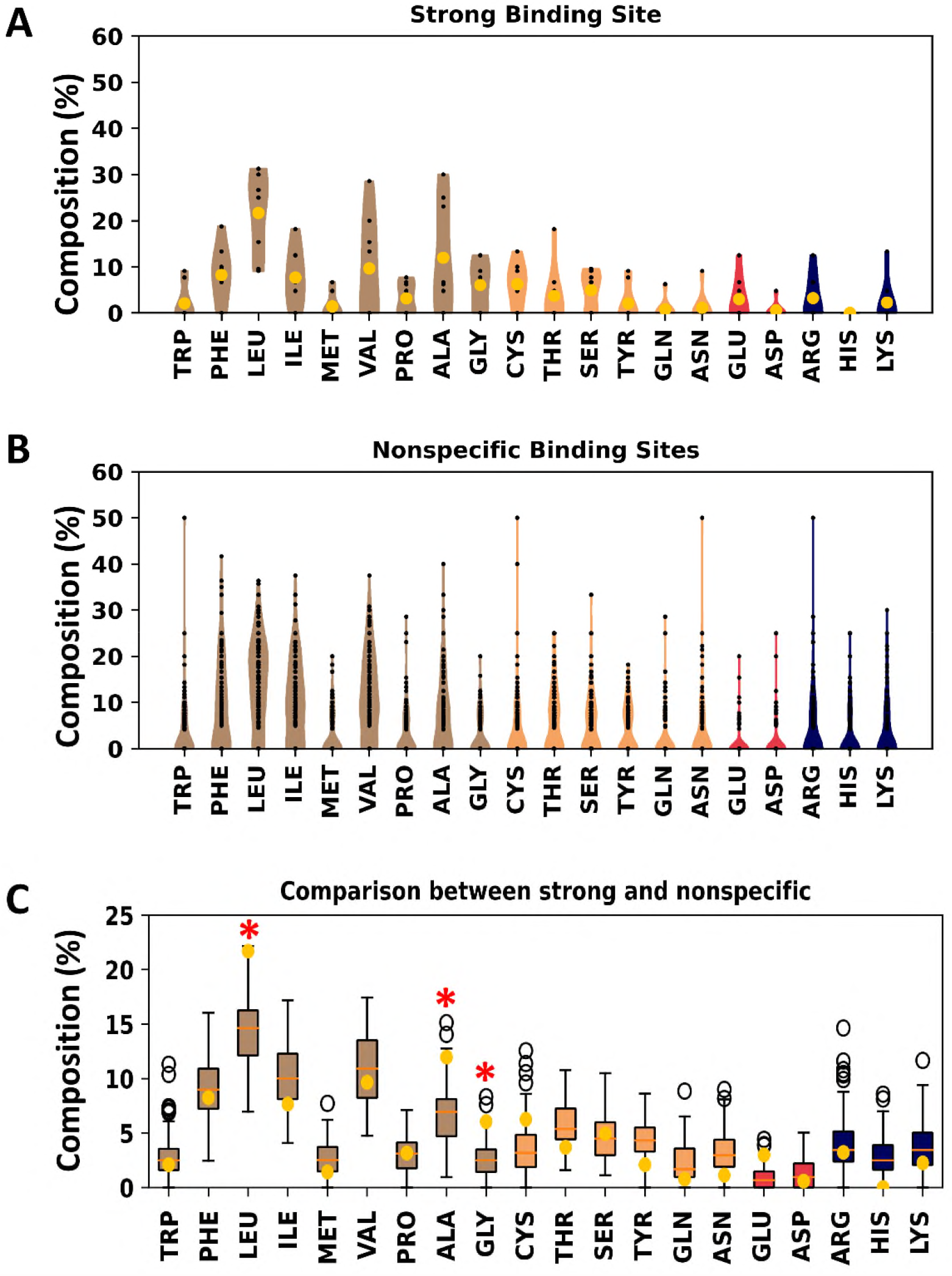
Amino acid composition of cholesterol binding site on GPCRs. (A) Violin plot of the amino acid composition of 8 specific binding sites that showed cholesterol residence times longer than 3 µs. (B) Violin plot of the amino acid composition of the 145 binding sites that showed shorter duration cholesterol interactions. (C) Comparison of the binding site amino acid compositions between the bootstrapping values from the 145 nonspecific binding sites (box plot) and the averages from the 8 specific binding sites (yellow dot). Data for amino acid compositions are color-coded based on the amino acid chemical property: data for non-polar amino acids are colored in brown, for polar amino acids in yellow, for acidic amino acids in red and for basic amino acids in blue. The red asterisks indicate residues where there is a clear difference in composition between the non-specific and specific site amino acid compositions.

We subsequently examined the representative cholesterol bound poses for these strong binding sites. These bound poses revealed two types of binding modes that are likely to have contributed to the stronger interactions in these binding sites. The first type features polar/charged interactions with the hydroxyl group of cholesterol, as seen in BS (binding site) id 5 of GCGR, and at BS id 7 and 4 of B2AR (Fig 6A and SI Fig S14). These polar/charged interactions may be the main stabilizing feature for strong cholesterol binding since the rest of the cholesterol molecule does not show extensive contacts with the receptor in these binding sites. The second type exhibits Leu sidechains at the rim of the sites that form a tight grip on the bound cholesterol molecule (Fig 6B and SI Fig S14). These residues might stabilize the cholesterol molecule in between the helices.

**Figure 6.**
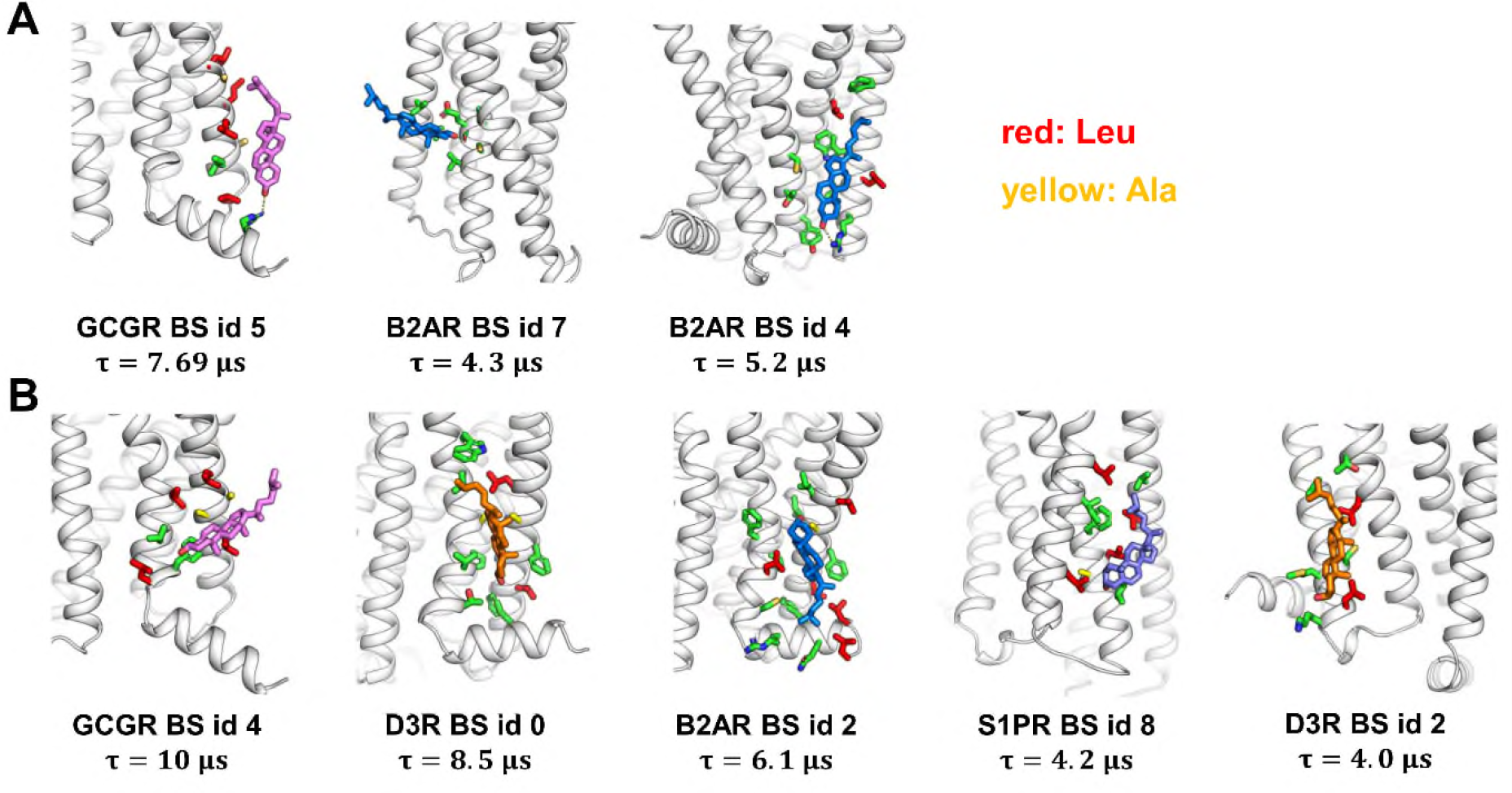
Representative cholesterol bound poses in the 8 specific binding sites. (A) Cholesterol bound poses with charge/polar interaction with the hydroxyl group. (B) Cholesterol bound poses without charge/polar interactions. Cholesterols are shown in sticks and colored based on the receptors they bound to. Protein residues within 0.5 nm of bound cholesterols are shown in green sticks. Text below each bound pose show the receptor name, the binding site (BS) id, and the calculated binding site residence time.

Taken together, PyLipID has allowed us to analyze cholesterol interactions efficiently and systematically with a set of 10 GPCRs. The analysis of 153 cholesterol binding sites revealed that most of cholesterols act as annular lipids around GPCRs, forming transient and potentially nonspecific interactions with the receptors. However, cholesterol may also form longer and more specific interactions with GPCRs at certain binding sites with distinctive structural features. The latter class of sites offer great potential as possible allosteric modulatory sites.

### Two examples of characterization of phospholipid interactions

We have also explored the application of PyLipID to interactions of membrane proteins with two (anionic) phospholipids, namely cardiolipin (for bacterial membrane proteins) and PIP_2_ (for mammalian membrane proteins). A recent survey of the energetics of membrane protein-lipid interactions as estimated by MD simulations ^51^ has shown that anionic phospholipids interact more strongly with membrane proteins (estimated free energies of -20 to -40 kJ/mol) than is the case for cholesterol (−5 to -10 kJ/mol). Thus, they are expected to exhibit longer residence times and provide good test cases for PyLipID analysis.

We have recently applied PyLipID to analyze cardiolipin interactions for a set of 42 *E. coli* inner membrane proteins based on CG-MD simulations using the Martini 3 force field ^52^. 700 cardiolipin binding sites were identified using PyLipID, analysis of which yielded a heuristic for defining a high affinity cardiolipin binding site, based on 2 or 3 basic residues in proximity, alongside the presence of at least one polar residue and one or more aromatic residues ^51^.

As an example of this analysis, we have selected formate dehydrogenase-N (PDB id 1KQF), a trimeric membrane protein, each subunit of which has five TM helices and a large cytoplasmic domain. The cardiolipin binding site observed in crystal structure was correctly identified by PyLipID as having the longest residence time among the 16 possible binding sites (Fig 7A and SI Fig S15). Analysis of residence times for individual binding site residues revealed that K254, K258, T39 and the main chain of P38 formed polar interactions or hydrogen bonds with cardiolipin headgroup, contributing to the main stabilizing force for the lipid bound poses in this binding site (Fig 7B). F37 also stabilized the bound lipid by hydrophobic stacking with the lipid tails.

**Figure 7.**
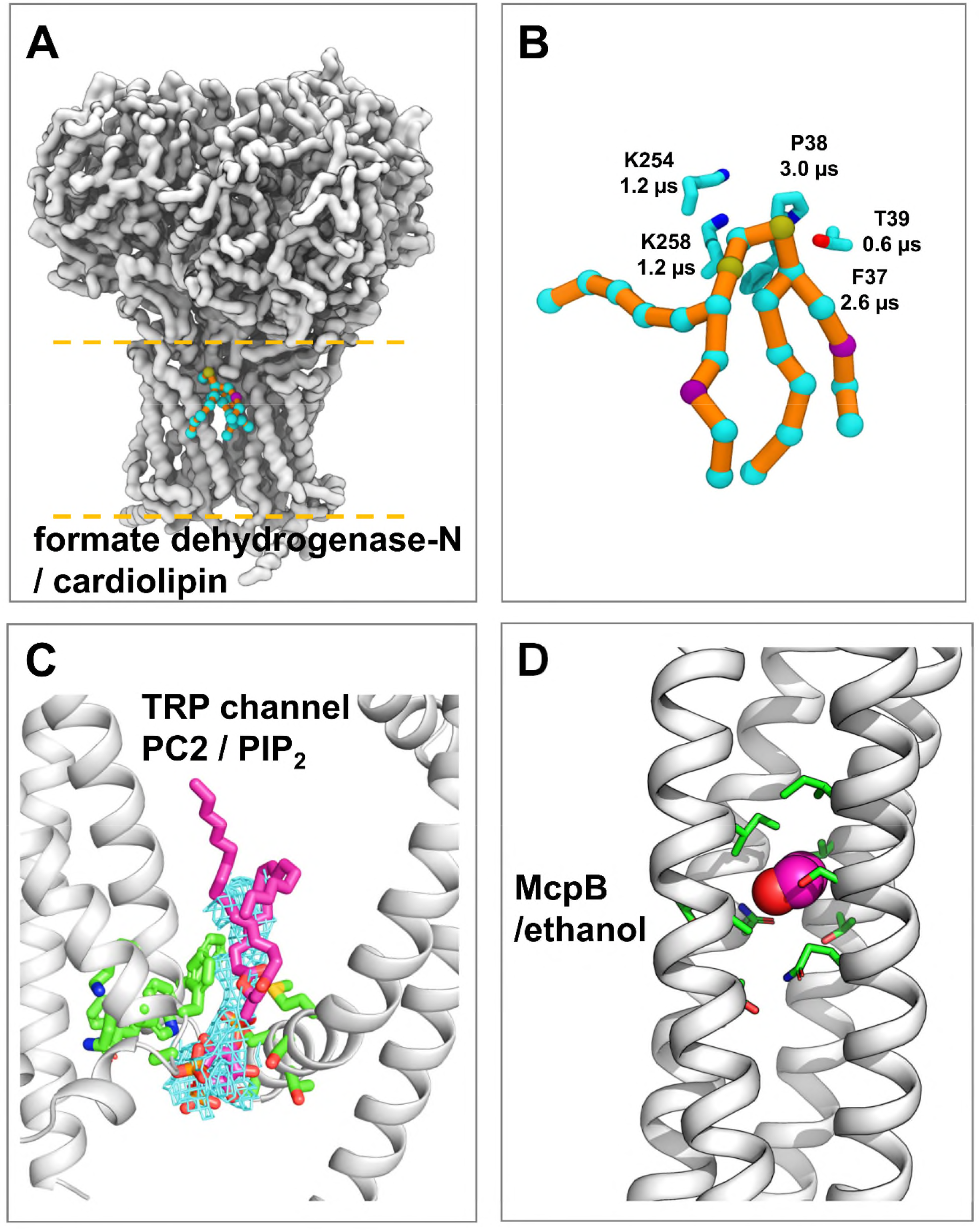
Application of PyLipID to phospholipids and non-lipid molecules. (**A**) Cardiolipin binding site with the longest residence time on formate dehydrogenase-N. The protein and lipid are described by the Martini CG model. The protein backbone beads are shown in white surface. The lipid beads are shown in cyan spheres connected by orange sticks. (**B**) A zoomed-in view of the cardiolipin binding site of formate dehydrogenase-N. The cardiolipin lipid is in the same representation as in panel A. Protein residues that showed the longest residence times in the binding site are shown in sticks. (**C**) The PyLipID calculated PIP_2_ binding site on the TRP channel PC2 overlaps well with the cryo-EM density. The PC2 cryo-EM structure is shown in white cartoon. The PIP_2_ density in the cryo-EM map is shown in blue mesh. The PIP_2_ bound pose calculated by PyLipID is shown in sticks in magenta. The binding site residues calculated by PyLipID are shown in sticks in green. This binding site showed the longest residence time in the Martini CG simulations, as calculated by PyLipID. **D** Ethanol binding sites on McpB. The main ethanol binding sites and an ethanol representative bound pose are shown. McpB is shown in white cartoon and ethanol in spheres. Key sidechains are in green.

In a second application of PyLipID to anionic lipids, we explored the interaction of PIP_2_ interaction with polycystin-2 (PC2), a TRP channel. A number of studies have implicated PIPs in TRP channel regulations ^12^. Based on CG-MD simulations in a membrane containing 10 % PIP_2_ in the cytoplasmic leaflet, 6 binding sites were identified from each of the four subunits of PC2 (SI Fig 6). The PIP_2_ binding site seen in the 3 Å resolution cryo-EM structure (PDB id 6T9N) ^53^ was identified by PyLipID as the site with the longest residence time. In addition, the representative bound pose of PIP_2_ in this binding site fits nicely within the lipid-like density in the cryo-EM map (Fig 7C). This again suggests that when multiple possible binding sites are present, residence time analysis using CG-MD simulations and PyLipID can be potentially used to identify the strongest interaction sites corresponding to lipid-like density observed by cryo-EM.

### Application to interactions of a non-lipid ligand with a membrane protein

PyLipID can be readily applied to characterize the binding non-lipid molecules in conjunction with atomistic simulations whenever sufficient binding/unbinding events are sampled. It therefore may be particularly useful for e.g. fragment screening approaches to binding site discovery (see ^54^ for an early application of this approach to GPCRs and ^55^ for a recent application using Martini 3).

To demonstrate the application of PyLipID to small-molecule/fragment binding, we analyzed the interactions of ethanol with a bacterial chemoreceptor, McpB, for which ethanol is a known attractant. The analyses were carried out on previously conducted atomistic simulations (3 × 600 ns) of an McpB cytoplasmic homodimer with 165 ethanol molecules (0.316 M) included to reproduce experimental conditions ^40^. As anticipated, ethanol molecules showed transient interactions with the receptor due to their small size and simple structure. Using PyLipID a total of 50 ethanol binding sites were identified on McpB, with residence times ranging from sub-nanosecond to ∼40 ns (SI Fig S16). Notably, the analysis highlighted several binding sites with longer residence times located within the center of the coiled-coil bundle (Fig 7D). It is suggested that these may facilitate conformational changes induced by ethanol binding to be transmitted to other parts of the receptor, thereby enabling the signaling response. To test the sensitivity of PyLipID to minor changes in protein sequence, we additionally analyzed atomistic simulations (3 × 600 ns) of McpB carrying the A431S mutation, which is known to considerably reduce taxis to alcohols ^40^. While the 51 ethanol binding sites identified by PyLipID largely overlap with those on wild-type McpB, ethanol binding to the sidechain of residue 431 was no longer observed (SI Fig S16). This example suggests therefore that PyLipID could be usefully employed as an analysis tool within an MD-based fragment screening study.

## DISCUSSION & CONCLUSIONS

### What does PyLipID allow us to do?

We have described PyLipID, an integrated package for analysis of protein-lipid interactions from MD simulation data. PyLipID has the following main features:

1. It calculates binding sites from simulation data using a robust methodology.
2. It calculates the residence times for lipid interactions with both the binding sites and individual amino acid residues.
3. It generates bound lipid poses and outputs structural representatives for each binding site.
4. It uses a dual-cutoff scheme to robustly quantify lipid interactions in a manner suitable for dynamic interactions in both coarse-grained and atomistic simulations.
5. It outputs interaction data in a convenient format to assist the ease and customization of subsequent large scale data analysis.

Thus, PyLipID provides for systematic and standardized analysis of protein-lipid interactions over large simulation datasets from multiple membrane proteins, facilitating comparative analysis of lipid binding sites. The inclusion of functions to generate representative bound poses allows for in-depth analysis alongside experimental structural data. PyLipID is an open-source Python package which allows users to customize the functions. It provides various portals for further manipulation of the generated data. It can be readily incorporated into analysis scripts, allowing for high through-put analysis of big data sets ^52^.

### How does PyLipID compare with other software in this area?

There are several frameworks developed for analysis of membrane MD simulations, building on the considerable expansion in this area of research over recent years. The closest in spirit to PyLipID is ProLint ^56^. ProLint is web-based, but also available as a standalone Python package *prolintpy*. ProLint provides feature-rich visualization and analysis tools, leaving binding site interpretations up to the user. In this respect it differs from PyLipID which automatically defines and analyses lipid binding sites to facilitate comparison with experiments, and to provide more directly pharmaceutically relevant structural insights. A somewhat simpler membrane protein simulation analysis framework is provided by MemProtMD ^57^, a database of CG-MD simulations of all known membrane protein structures in a model bilayer, which provides contact-based metrics for protein-lipid interactions, and information local bilayer thickness distortion by proteins. MemProtMD is now directly linked to membrane protein entries by the RCSB/PDB. There have also been several recent packages developed which are aimed at analysis of lipid bilayers. These include e.g. LiPyphilic ^58^, which is a fast Python package for analyzing complex lipid bilayer simulations (but not yet extended to membrane proteins), and FATSLiM ^59^, also in Python, which enables bilayer leaflet identification, bilayer thickness and area per lipid calculations, and which works for various (curved) membrane geometries and bilayers including proteins. In terms of more detailed analysis of interactions at binding sites, there are several more general approaches for drug-target residence times via simulations, including e.g. τRAMD ^60^ which may in principle be adaptable to protein-lipid interactions.

### What can PyLipID teach us about protein lipid interactions?

We have described a couple of applications of PyLipID. There is a considerable literature on identifying and characterizing GPCR/cholesterol interactions by MD simulations (e.g. ^61-63^) and it is not our aim to review these here (for recent reviews see e.g. ^12, 64^). There have also been a number of GPCR structural studies e.g. combined with docking of cholesterol ^65^ to generate a database of predicted binding sites for cholesterol on membrane proteins, or via analysis of crystal structures of GPCRs with bound cholesterol molecules ^49^. PyLipID provides some new insights into GPCR/cholesterol interactions. In particular, the analysis of residence times has allowed us to separate interactions/sites in annular and specific cholesterol binding sites, the latter showing longer residence times and with enriched interactions with Leu, Gly and Ala residues. Extending this approach to a couple of anionic phospholipids suggests that long residence time binding sites correlate with those observed experimentally in cryo-EM structures, indicating how PyLipID may be used to aid the assignment and analysis of lipid-like density in newly determined structures ^66^.

## ACKNOWLEDGEMENTS

We thank Yi Hao for designing the PyLipID logo. Research in the M.S.P.S. group is supported by Wellcome (208361/Z/17/Z), BBSRC (BB/R00126X/1), and PRACE (Partnership for Advanced Computing in Europe; 2016163984). Research in P.J.S. lab is supported by Wellcome (208361/Z/17/Z), BBSRC (BB/P01948X/1, BB/R002517/1 and BB/S003339/1) and MRC (MR/S009213/1). W.S. acknowledges support from the Newton International Fellowship. T.B.A. is supported by a Wellcome studentship (102164/Z/13/Z). This project made use of time on ARCHER via the HECBioSim consortium, supported by EPSRC (EP/L000253/1).

