## Supplementary material for "PyLipID: A Python package for analysis of protein-lipid interactions from MD simulations": SI

**SI FIGURES**

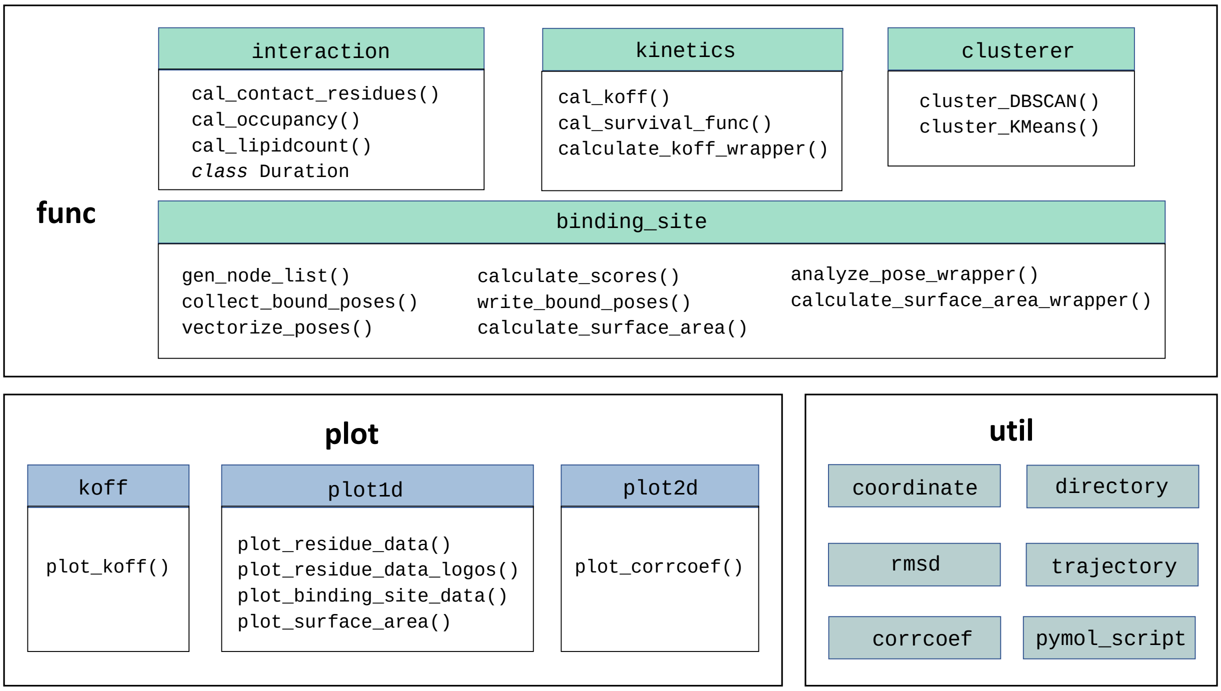


**SI Figure 1. PyLipID package design: `func`, `plot` and `util` modules.** These three modules are the inner layer of the package that contain functions for doing the heavy-lifting. The `func` module includes functions for calculation of lipid interactions, and can be divided into four sub-modules: `interaction` that contains functions/classes for calculation of durations, occupancy and lipid count; `kinetics` that contains functions for calculation residence time/$k_{off}$; `clusterer` that contains functions to cluster the bound poses; and `binding_site` that contains functions for calculation of binding site related properties. The `plot` module provides the assisting functions for plotting, and can be divided into three sub-modules: `koff` that contains the function for plotting $k_{off}$s; `plot1d` that contains functions for plotting properties as a function of residue index or binding site index; and `plot2d` that contains the function for plotting correlation. The `util` module provides various utility functions including creating directories, saving coordinates, and defining functions independent of PyLipID workflow.

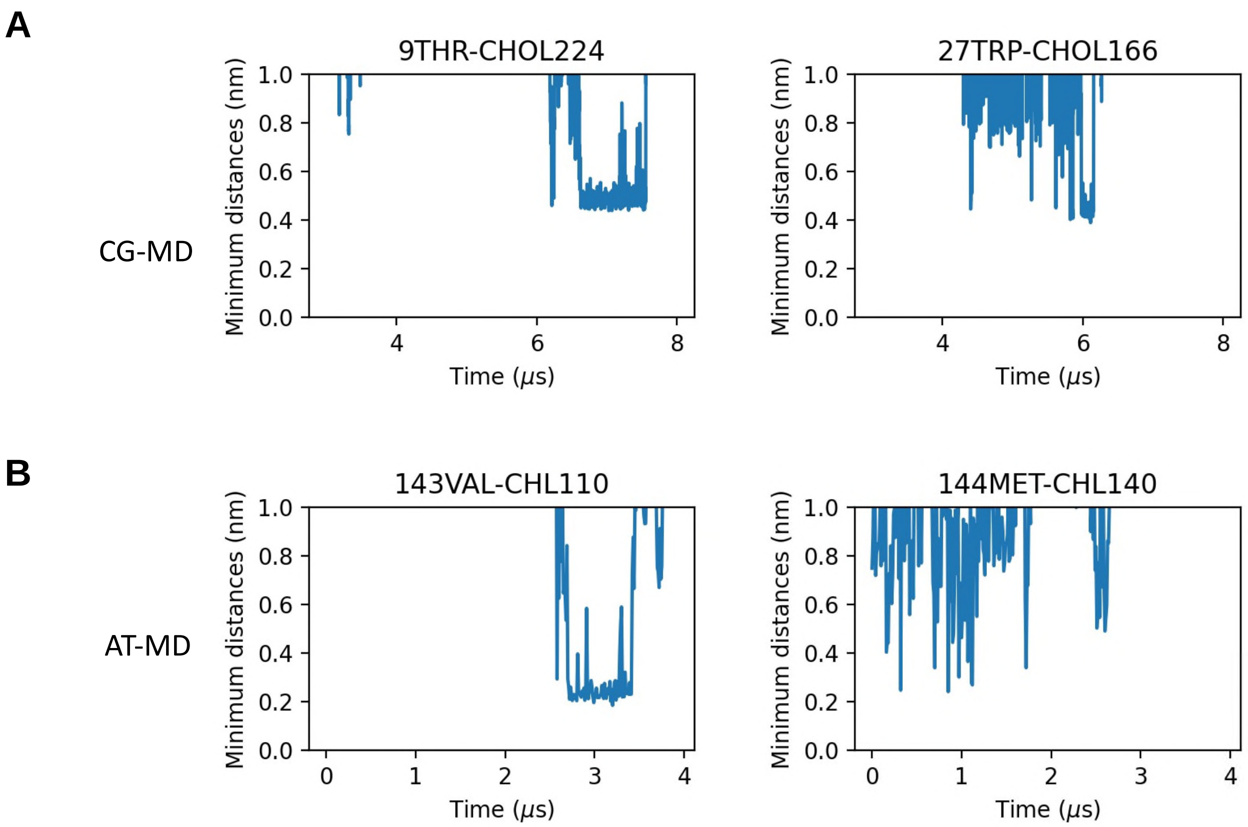


**SI Figure S2.** Minimum distances of cholesterol molecules to their contacting residues in (A) the coarse-grained simulation of A2a receptor and (B) in the atomistic simulation of GCGR. The left column illustrates the 'rattling in the cage' effect for contacting molecules, whereas the right column shows the minimum distances of non-contacting molecules. The distance of wiggling molecules can often overlap with the distances of the non-contacting molecules.

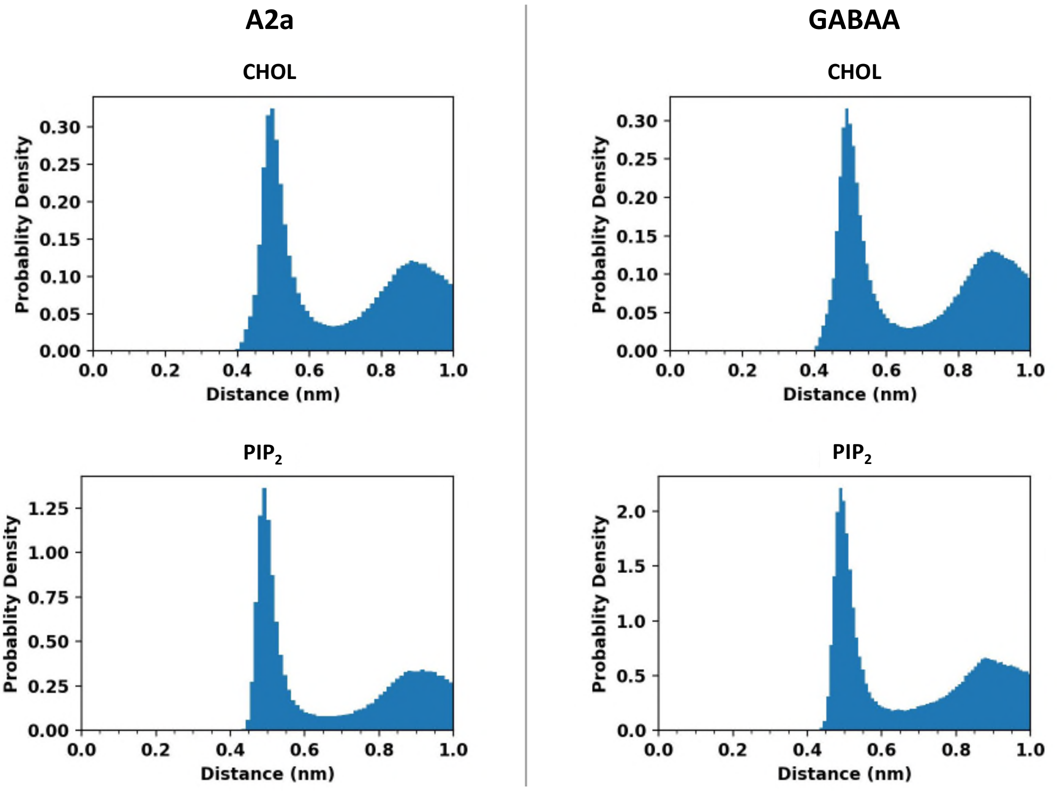


**SI Figure S3.** Distribution of the minimum distances of cholesterol and $PIP_2$ molecules in the coarse-grained simulations of A2a receptor and GABAA receptor. The same lipid species showed similar distance distributions in the two systems. However, the two lipid molecules have different peaks and different starting points of their distance distributions.

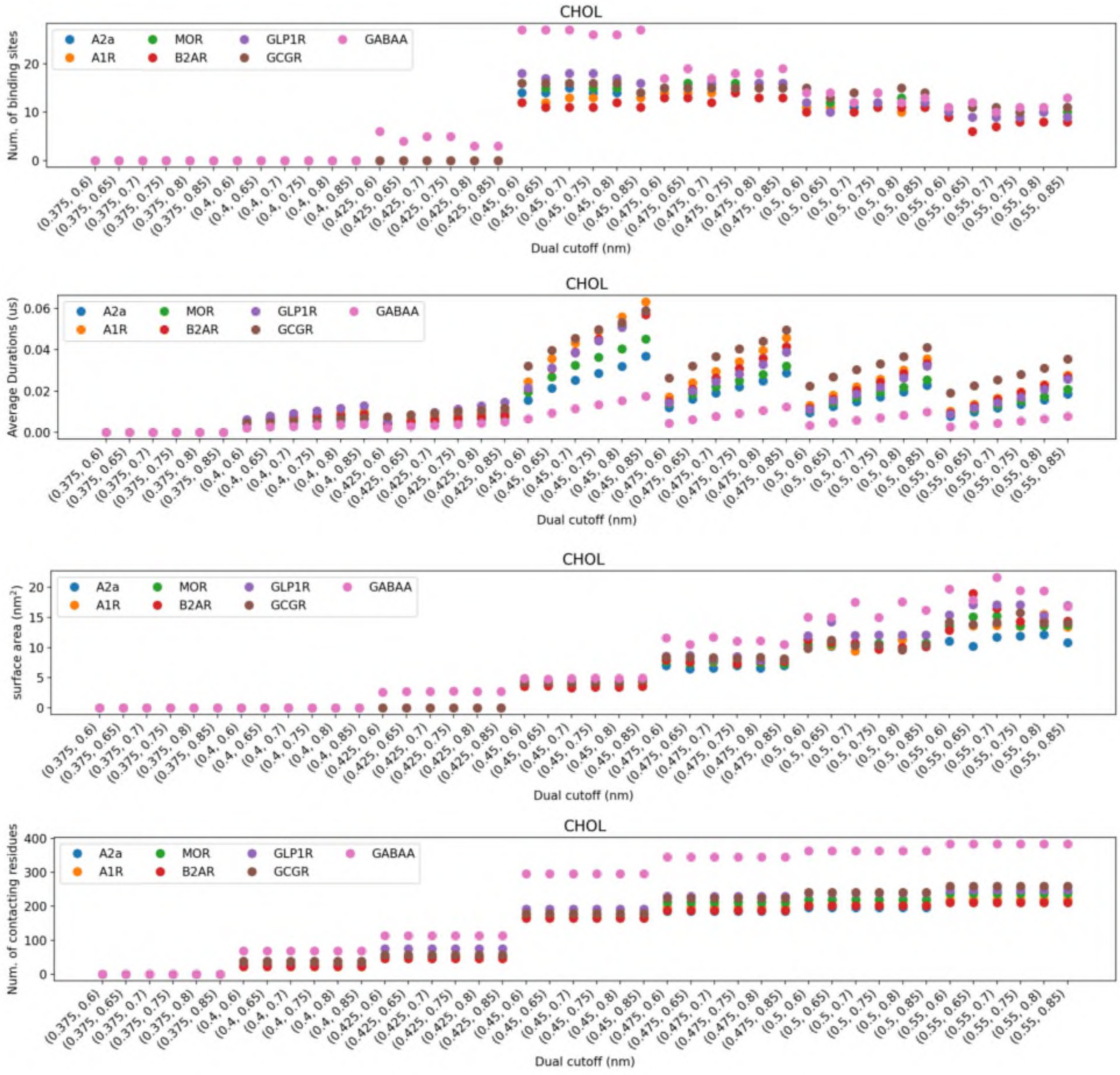


**SI Figure S4.** Cholesterol interactions calculated by PyLipID using different dual cut-off values. The number of binding sites, the average contact durations, and the average surface area of binding sites of cholesterol interactions were calculated from the coarse-grained simulations of A2aR, A1R, B2AR, μOR, GLP1R, GCGR and GABAA using different dual cut-offs.

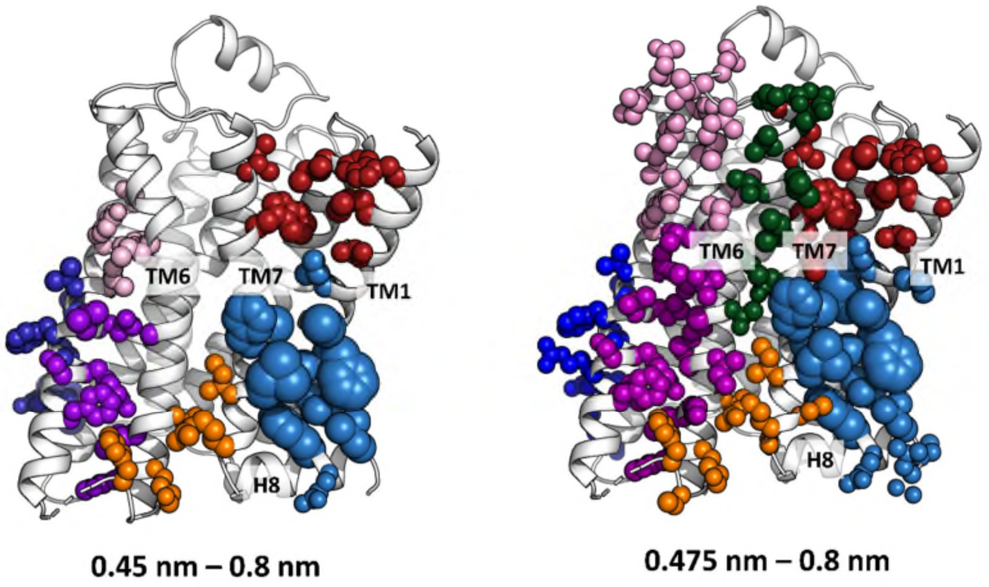


**SI Figure S5**. Comparison of cholesterol binding sites from using different lower cut-off values. The cholesterol binding sites on A2aR were calculated using two dual-cutoffs, i.e. 0.45 nm - 0.8 nm and 0.475 nm - 0.8 nm. The calculated binding sites on the side of TM5_TM6_TM7 are shown, and residues comprising the same binding sites are shown in the same colour in both results. Binding site residues are shown in spheres with their spheres corresponding to their residence time of cholesterol interactions.

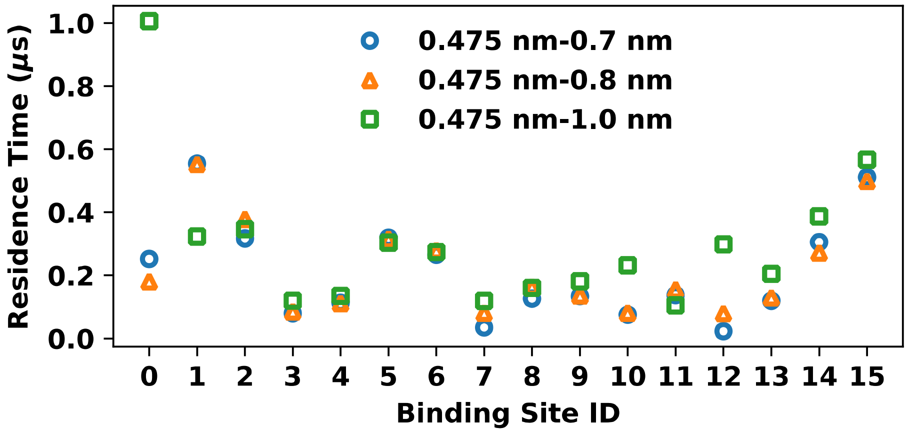


**SI Figure S6.** Comparison of binding site residence times from using different upper cut-off values. The residence times of cholesterol binding sites on A2aR were calculated using three dual-cutoffs, i.e. 0.475-0.7, 0.475-0.8 and 0.475-1.0. The three dual-cutoffs identified the same cholesterol binding sites on A2aR. The binding site residence times calculated from 0.475-0.7 and 0.475-0.8 were very similar, whereas those from 0.475-1.0 showed some differences at a couple of binding sites.

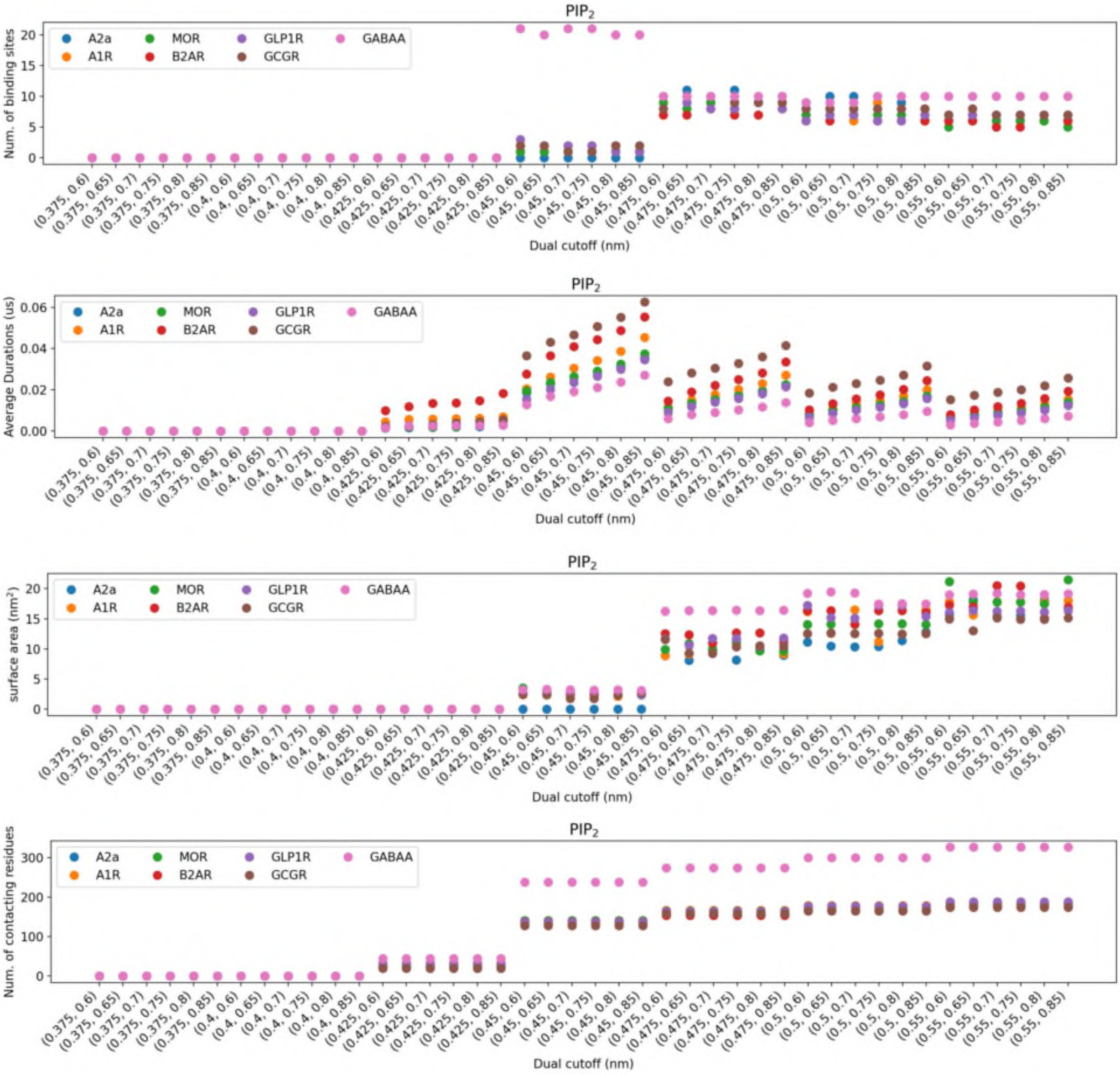


**SI Figure S7.** $PIP_2$ interactions calculated by PyLipID using different dual cut-off values. The number of binding sites, the average contact durations and the average surface area of binding sites of cholesterol interactions were calculated from the coarse-grained simulations of A2aR, A1R, B2AR, µOR, GLP1R, GCGR and GABAA using different dual cut-offs.

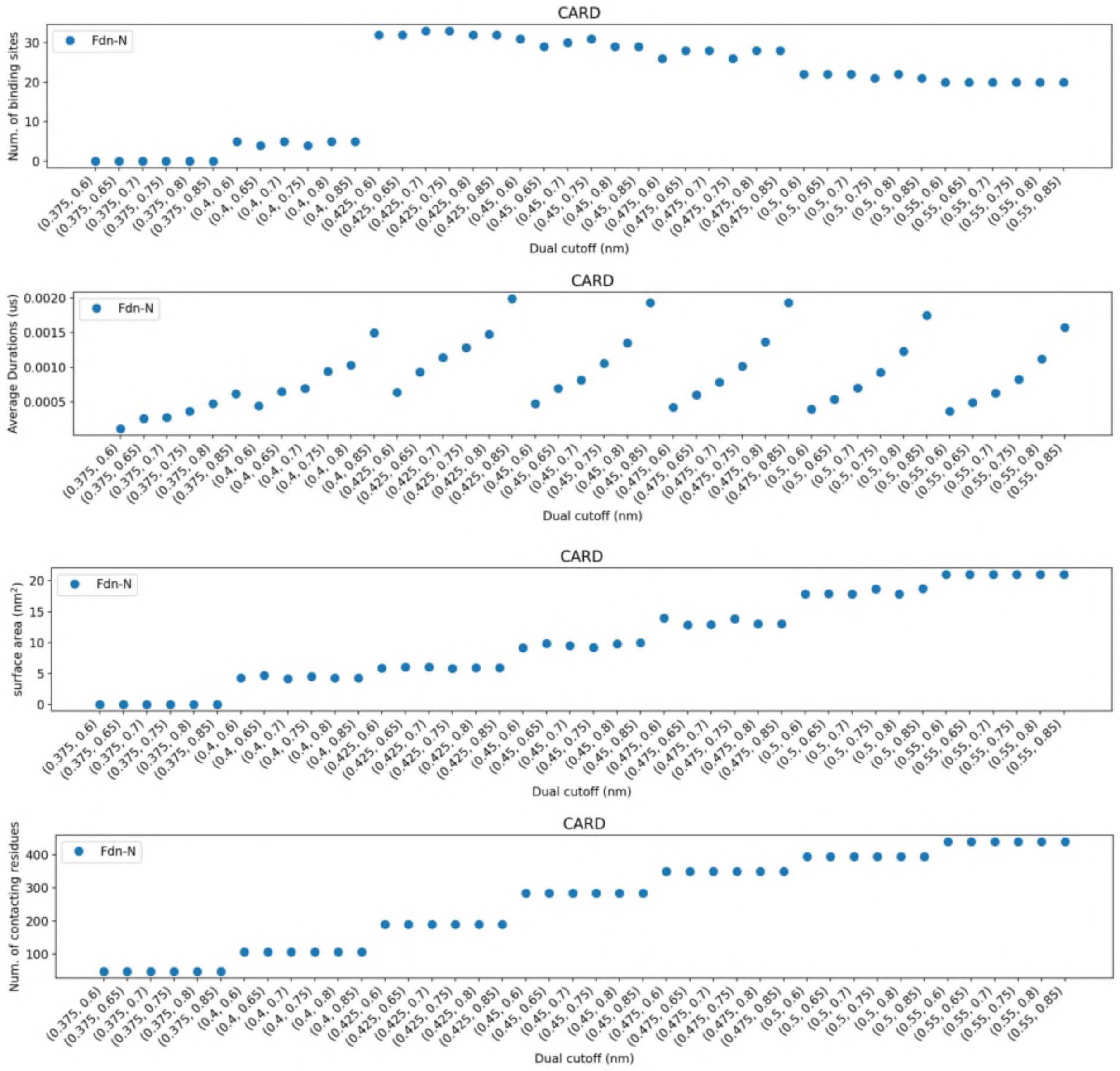


**SI Figure S8.** Cardiolipin interactions with formate dehydrogenase-N (Fdn-N) using a sequence of dual cut-offs.

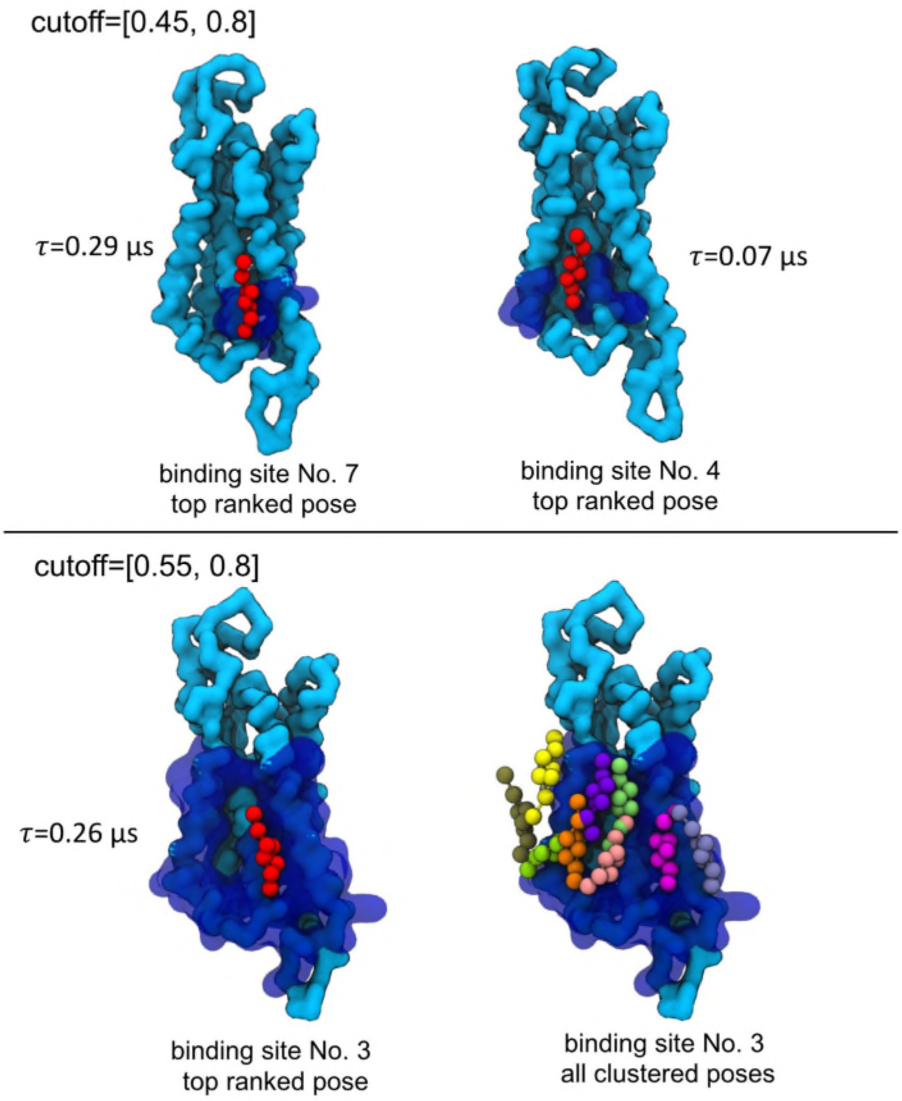


**SI Figure S9.** Comparison of the representative bound poses generated by the dual cut-off values of 0.475 nm - 0.8 nm and 0.55 nm - 0.8 nm.

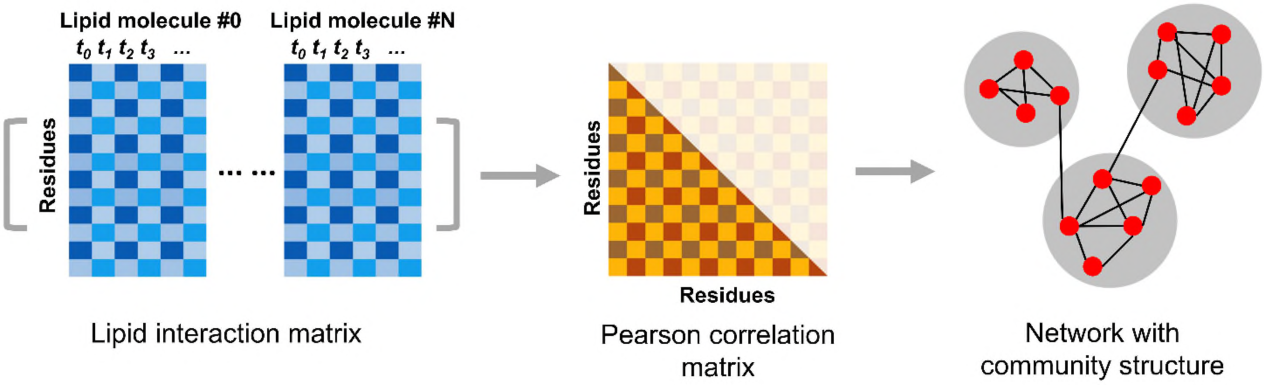


**SI Figure S10. Calculation of binding site from interaction network**. The interactions of each protein residue with the lipid molecules are recorded as a function of time to build a lipid interaction matrix. The Pearson correlation of the residues is then calculated from this lipid interaction matrix, and then used to build a lipid interaction network in which the nodes are protein residues, and the edges are the interaction correlation between the two connecting residues. The binding sites are calculated as the community structures of this interaction network.

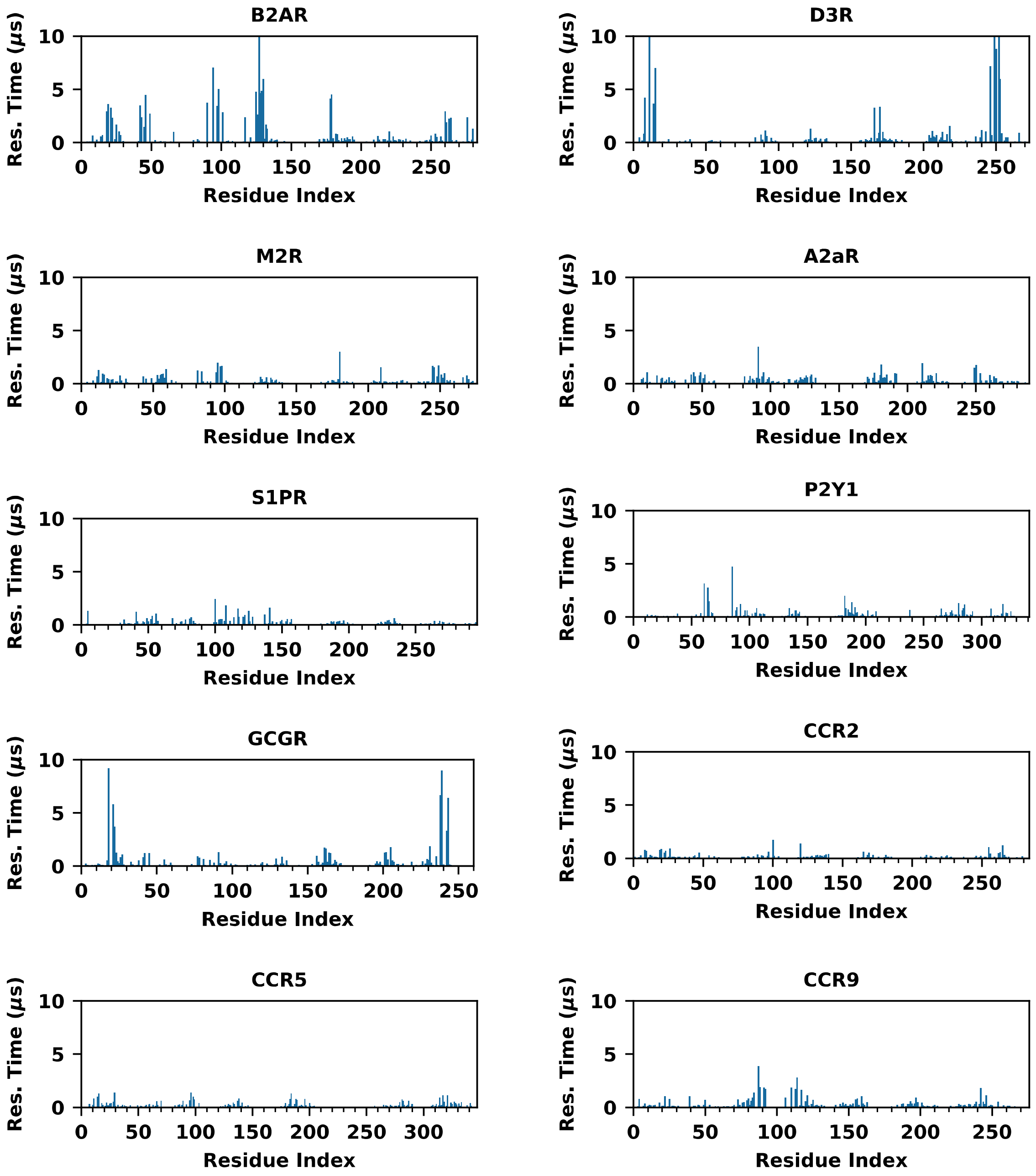


**SI Figure S11. Residence time of cholesterol interaction on GPCRs as a function of protein residue index.**

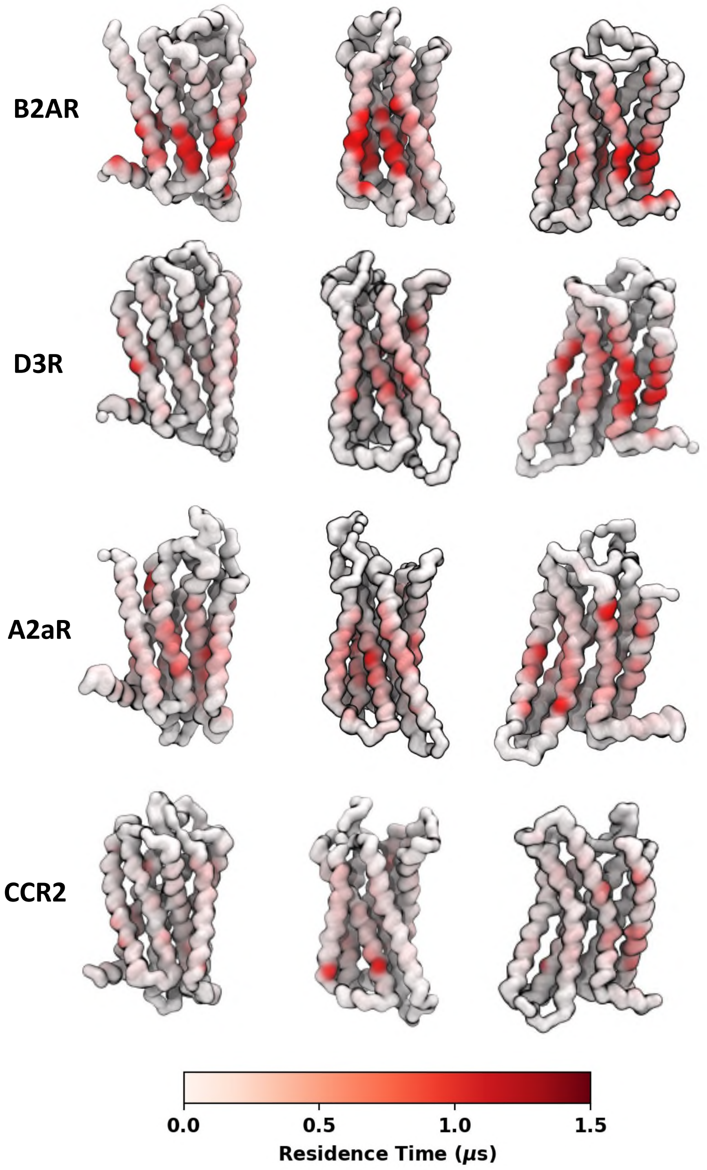


**SI Figure S12. Hotspots of cholesterol interactions on GPCRs**. B2AR showed strongest cholesterol interactions; D3R had strong interactions at a couple of location including between TM1 and TM7; A2aR showed intermediate level of interactions with cholesterols; CCR2 showed weak and non-specific interactions with cholesterols. The receptors are in MARTINI coarse-grained models. Only backbone beads are shown to make the secondary structure clear. The surfaces of the proteins are colored white-to-red based on the residue cholesterol residence times.

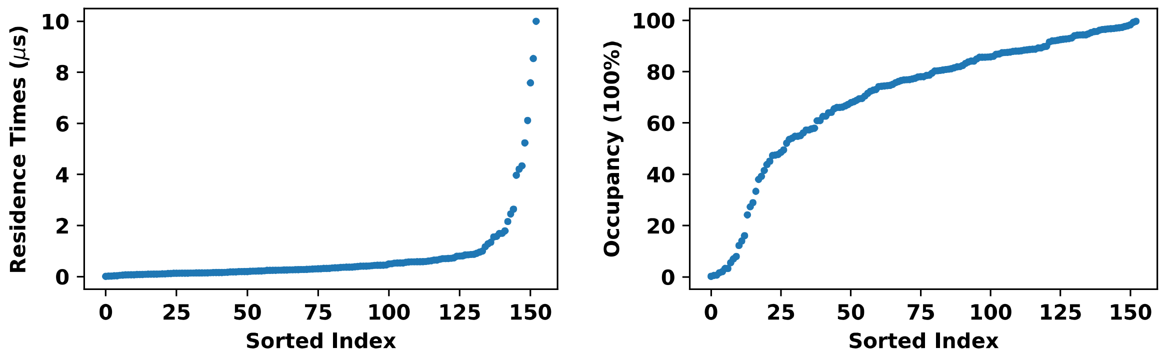


**SI Figure S13. Ranking of cholesterol binding sites on GPCRs.** (Left) Plot of binding site residence times in a sorted order. (Right) Plot of binding site occupancies in a sorted order.

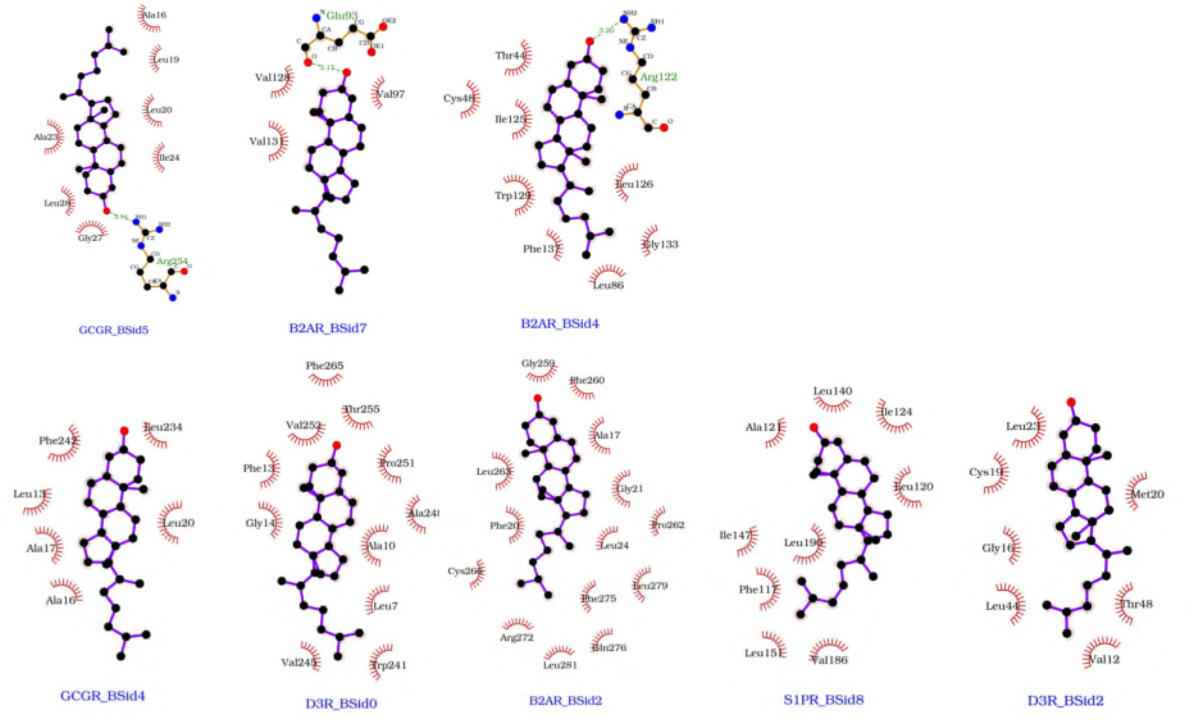


**SI Figure S14. Cholesterol binding site on GPCRs.** The schematic presentation of cholesterol interactions is generated for the 8 binding sites that showed cholesterol residence time longer than 3 μs. Similar to Fig 8, the upper row includes binding sites in which cholesterol had charged/polar interactions on its hydroxyl group; and the lower row includes binding sites that contain only non-polar/hydrophobic interactions. The schematic plots were generated using LigPlot+ [1], and the representative bound pose of the binding site.

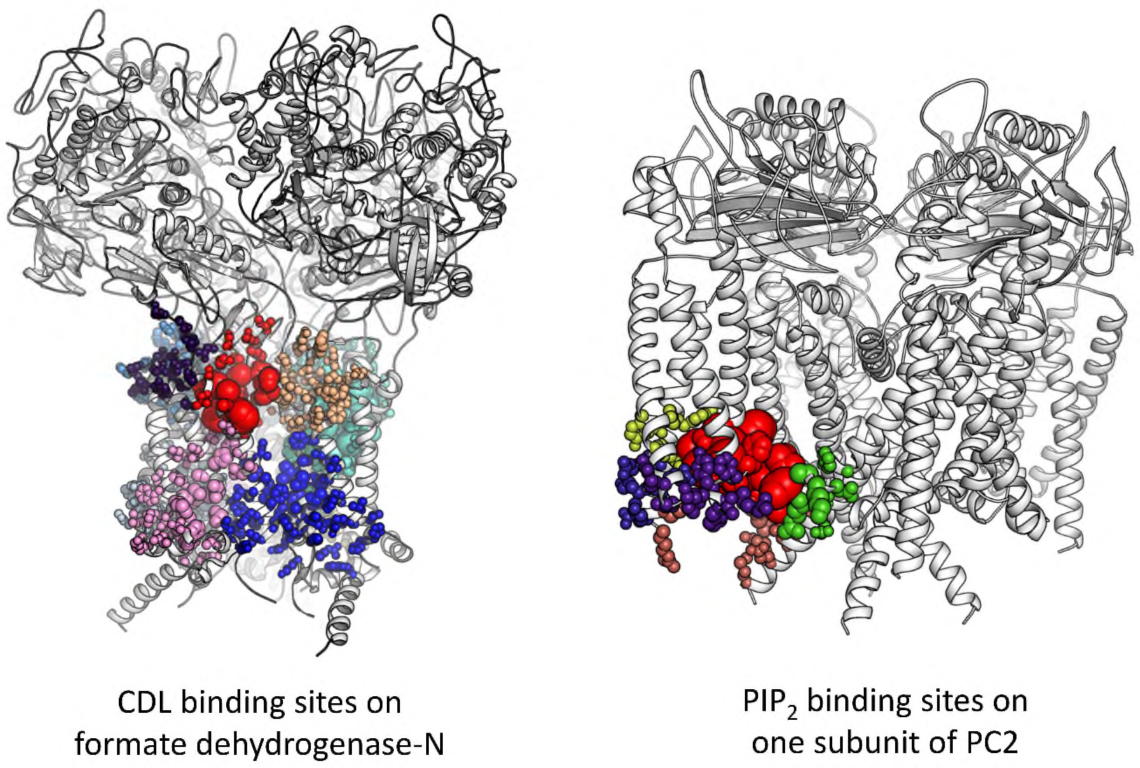


**SI Figure S15. Cardiolipin and PIP$_2$ binding sites**. (Left) The calculated cardiolipin binding sites on formate dehydrogenase-N by PyLipID were mapped onto the receptor structure. (Right) The calculated binding site of PIP$_2$ headgroups were mapped onto one of the four identical subunits of PC2. In both cases, proteins are shown white cartoon and binding site residues are shown in spheres. Residues from the same binding site are shown in the same colors, and the sphere scales of binding site residues corresponding to the residue residence times. The binding sites revealed by the experimental structures are colored red. The presentations are made by PyMol session, which is generated by a python script created by the method of `save_pymol_script()` of PyLipID.

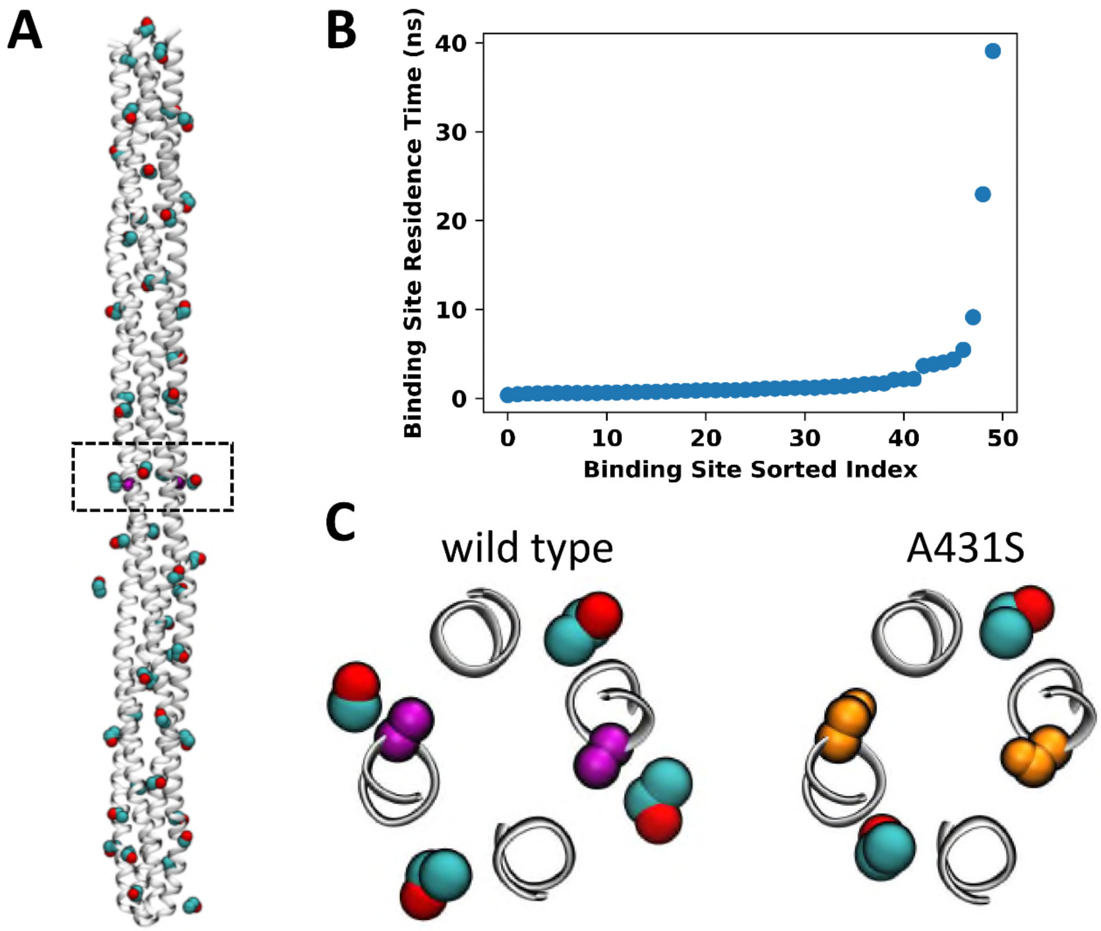


**SI Figure S16. Ethanol binding sites on McpB.** (A) Ethanol binding sites and representative bound poses on wild type McpB. McpB is shown in white cartoon with ethanol shown as colored spheres. The dashed box highlights ethanol binding near residue A431 (purple spheres) (B) Sorted residence times of ethanol binding sites on wild type McpB. (C) Comparison of identified ethanol binding sites for wild-type and A431S McpB. Residue 431 is shown as purple (alanine) or orange (serine).

**SI TEXT**

**Choice of cut-offs and its impact on the reported values.**

PyLipID defines contacts based on the measure of minimum distances, thus the accuracy of the calculation of lipid interactions, especially the calculation of binding site, will be determined by the choice of the distance cut-offs. By default, PyLipID uses a dual cut-off scheme in which a continuous contact starts when the minimum distance to the target residue is smaller than the lower cut-off and ends when the minimum distance is larger than the upper cut-off. This scheme was created to deal with the 'rattling in the cage' effect in coarse-grained simulations, in which molecules undertake rapid motions at their site without real dissociation (SI Fig S2). The contact distance and cutoff distances may vary among different lipid species. Comparison of the probability densities of cholesterol and $PIP_2$ molecules from the A2aR and GABAAR (PDB id 6HUP) simulations revealed that $PIP_2$ contact starts ~0.44 nm and the first peak of annular lipids locates at 0.49-0.5 nm, whereas cholesterol contact start at ~0.39 nm and the first peak of annular lipids at ~0.5-0.51 nm (SI Fig S3).

Based on how contacts are calculated in PyLipID, it is anticipated that the lower cut-off will affect the detection of lipid contacts and lipid binding sites, whereas the upper cut-off will affect the durations of these contacts. To illustrate the impact of cut-offs, we plotted the number of binding sites, the average durations of contacts, the average binding site surface area and the number of contacting residues for a sequence of dual cut-offs on a couple of GPCRs, including A2aR, A1R, µOR, ß2AR, GCGR and GLP1R, and a ligand-gate ion channel GABAA (SI Fig S4). The lower cut-off ranged from 0.3 nm to 0.55 nm and the upper cut-off from 0.6 nm to 0.85 nm. For cholesterol interactions, GABAA and GPCRs showed different changes of e.g. the number of binding sites, the averaged interaction durations, the number of contacting residues, in response to different cut-off values, due to the different protein structures and chemical properties of the protein surfaces of these two receptor families. For example, cholesterol has shorter contact durations with GABAA regardless of the cut-off values, and more calculated binding sites for the lower cut-offs of 0.45 nm and 0.475nm, compared to the tested GPCRs. The diverse profiles of the two protein families could help to find some consistent rules concerning the choice of cut-off values.

Let us first look at the lower cut-off. Cholesterol contacts started to be detected at the lower cut-off of 0.4 nm in both protein families, whereas cholesterol binding sites were detected at 0.45 nm in GPCRs and at 0.425 nm in GABAA. The starting distance for binding site detection is governed by the structure and geometry of the transmembrane region. The shorter lower cut-off for detection of binding sites in the latter system is due to that GABAA has straight transmembrane helices and the cholesterol binding sites on this protein largely consist of two adjacent helices and most on the surface of the protein. In comparison, GPCRs have more complex structures in the transmembrane region with crooked and tilted helices and the binding sites often span cross multiple helices and can reach deeper into the helix clefts. The binding sites detected with the minimal lower cut-off, i.e. 0.45 nm in GPCRs and 0.425 nm in GABAA, were very fragmented, consisted of three to four residues, as indicated by their small surface areas. This is because that this small cut-off can only detect the synergistic binding activities of residues that showed the closet contacts with the lipid molecules. Whereas the use of this small lower cut-off may reveal some critical residues that showed strongest correlation of binding lipids, it can miss many important binding information. As illustrated in the comparison of cholesterol binding sites in B2AR using two dual-cutoffs, i.e. 0.45 nm – 0.8 nm and 0.475 nm – 0.8 nm (SI Fig S5), some of the binding sites that comprised of helices further apart from each other (i.e. the green binding site between TM6 and TM7 in SI Fig S5) cannot be detected by the lower cut-off of 0.45 nm. In addition, the larger lower cut-off of 0.475 nm also gave a fuller list of residues for each binding site which provides a better description of the binding sites.

In both protein systems, the number of binding sites started to decrease at the lower cut-off that is 0.5 nm larger than the minimal lower cut-off for binding site calculation, i.e. 0.5 nm in GPCRs and 0.475 nm in GABAA. This suggests that too large lower cut-offs can merge binding sites and lower the resolution of reported results. *Therefore, a general rule for determining the value for the lower cut-off is to take the value right before the number of binding sites starting to decrease, i.e. 0.475 nm for GPCRs and 0.45 nm for GABAA.*

The upper cut-off mostly affected the interaction durations, as shown in the increase of averaged interaction durations with the increase of upper cut-off. Due to the dynamical behaviour of protein conformations, the interaction network can fluctuate with different interaction durations, i.e. different upper cut-offs. Thus, we saw fluctuations of the number of detected binding sites and the surface areas when the lower cut-off was fixed but the upper cut-off increased from 0.6 nm to 0.85 nm. *Based on our observations, the upper cutoffs ranging from 0.7 nm to 0.85 nm gave quite consistent results for binding site calculation, whereas too small or too large a upper cutoff can generate deviations* (SI Fig S6).

The effective cut-off values can vary for different lipid species. For $PIP_2$ interactions on the same set of simulations systems, the lipid contacts started to appear with the lower cut-off of 0.425 nm and the lipid binding sites at 0.45 nm (SI Fig S7). This agrees with the distribution of minimum distances which started at a larger distance than that of cholesterol. The number of binding sites and the binding site surface areas also showed larger fluctuations with different upper cut-off, largely due to the high dynamics of lipid tails*. For PIP2 calculations, we suggest that the lower cut-off in the range of 0.5 nm to 0.6 nm normally work well.* We also checked the cardiolipin interactions with the *E coli* protein formate dehydrogenase-N (Fdn-N) (SI Fig S8). The dependence on cut-offs again different from the other two lipid species.

**Choice of cut-offs and its impact on the representative bound poses.**

The representative bound poses, i.e. bound pose with the largest samplings from simulations, can always be picked up regardless of the choice of cut-off values if these values can effectively detect binding sites. This is because PyLipID uses a density-based scoring function to value bound poses, so that frequently sampled poses always rank high regardless of the binding site definition. The cut-off values, however, affects how many representative bound poses being generated via affecting the number of binding sites being detected. For example, in the case of using 0.45 nm - 0.8 nm cut-offs, PyLipID detected two cholesterol binding sites around ICL2 of A2aR, one is between TM3 and TM5 and the other between TM3 and TM4, but the former has longer interaction residence time and more samplings. The representative bound poses for these two binding sites were generated by PyLipID and shown in SI Fig S9. When we used a larger lower cut-off, i.e. 0.55 nm - 0.8 nm, these two binding sites were merged into one and the representative bound pose was the one between TM3 and TM5. The possible bound poses in a binding site can be checked by clustering the bound poses, and the clustered results can reveal the less frequently sampled bound poses. In this case, when we checked the all the clustered bound poses, the one between TM3 and TM4 was revealed. *Thus, too big a lower cut-off may hide some of the interesting poses, whereas small lower cut-offs may generate unwanted noises and also lose some of the geometry features of binding sites.*

**SI TABLES**

**SI Table S1 Simulation overview***

| ***GPCRs*** | | | |
|---|---|---|---|
| MARTINI coarse-grained simulations | | | |
| Receptor | PDB ID | Length x repeats | Membrane lipid composition |
| B2AR | 2RH1 | 10 µs x 3 repeats | Upper leaflet: POPC(65%):CHOL(35%)<br>Lower leaflet: POPC(55%):CHOL(35):$PIP_2$(10%) |
| D3R | 3PBL | | |
| M2R | 4MQT | | |
| A2aR | 3EML | | |
| S1PR | 3V2Y | | |
| P2Y1 | 4XNV | | |
| GCGR | 5EE7 | | |
| CCR2 | 5T1A | | |
| CCR5 | 4MBS | | |
| CCR9 | 5LWE | | |
| ***Polycystin-2*** | | | |
| MARTINI coarse-grained simulations | | | |
| Polycystin-2 | 6T9N | 100 µs x 3 repeats | Upper leaflet: POPC (100%)<br>Lower leaflet: POPC(90%):$PIP_2$(10%) |

* For simulation details of the formate dehydrogenase-N system, see [2], and details of the McpB system, see [3].

### SI METHODS

#### System setup

For the 10 GPCR simulations, the clean and processed receptor coordinates were downloaded from MemProtMD database [4] (http://memprotmd.bioch.ox.ac.uk). The protein coordinates were converted into MARTINI coarse-grained models using *martinize.py*[5], and then embedded in an asymmetrical membrane using *insane.py*[6] with a lipid composition shown in SI Table 1. Systems were solubilized with Martini waters and ions to a neutral charge. Systems were minimized using the steepest descent method, then equilibrated for 100 ns. The equilibration step used a semi-isotropic Berendsen barostat[7] at 1 bar, and a velocity-rescaling thermostat[8] at 310 K. Production simulations were then run using the Parrinello-Rahman barostat[9] at 1 bar using 20 fs. These simulations were run using Gromacs 2020[10].

For the PC2 simulations, the structure was obtained from the PDB (PDB id 6T9N) and missing loops modelled using MODELLER 9.20[11]. The structure was converted to coarse-grained resolution using the MARTINI 2.2 forcefield and *martinize.py*. The ElNeDyn elastic network with a cut-off of 0.9 nm and a force constant of 1000 kJ $mol^{-1}$ $nm^{-2}$ was applied. The protein was embedded into the bilayer and solvated with MARTINI water using *insane.py*. NaCl was added to an approximate concentration of 0.15 M. The system was minimized via the steepest descent method and equilibrated in 2x 100 ns steps. 3x 100 µs CG simulations were performed using a 20 fs timestep. The temperature was maintained at 310K using the V-rescale thermostat and the pressure maintained at 1 bar using the Parrinello-Rahman barostat. Simulations were run using Gromacs 2019.

For simulation details of the formaet dehydrogenase-N system, see [2], and details of the McpB system, see [3].

#### Data Analysis

To assist structural analysis, the bound poses from coarse-grained simulations were converted to atomistic structures using CG2AT2[12]. Images were made with VMD[13] and PyMol[14]. Plots made with Matplotlib[15].
